# A comprehensive analysis of the global human gut archaeome from a thousand genome catalogue

**DOI:** 10.1101/2020.11.21.392621

**Authors:** Cynthia Maria Chibani, Alexander Mahnert, Guillaume Borrel, Alexandre Almeida, Almut Werner, Jean-François Brugère, Simonetta Gribaldo, Robert D. Finn, Ruth A. Schmitz, Christine Moissl-Eichinger

## Abstract

The human gut microbiome plays an important role in health and disease, but the archaeal diversity therein remains largely unexplored. Here we report the pioneering analysis of 1,167 non-redundant archaeal genomes recovered from human gastrointestinal tract microbiomes across countries and populations. We identified three novel genera and 15 novel species including 52 previously unknown archaeal strains. Based on distinct genomic features, we warrant the split of the *Methanobrevibacter smithii* clade into two separate species, with one represented by the novel Candidatus *M. intestini*.

Patterns derived from 1.8 million proteins and 28,851 protein clusters coded in these genomes showed substantial correlation with socio-demographic characteristics such as age and lifestyle. We infer that archaea are actively replicating in the human gastrointestinal tract and are characterized by specific genomic and functional adaptations to the host. We further demonstrate that the human gut archaeome carries a complex virome, with some viral species showing unexpected host flexibility. Our work furthers our current understanding of the human archaeome, and provides a large genome catalogue for future analyses to decipher its role and impact on human physiology.

GRAPHICAL ABSTRACT

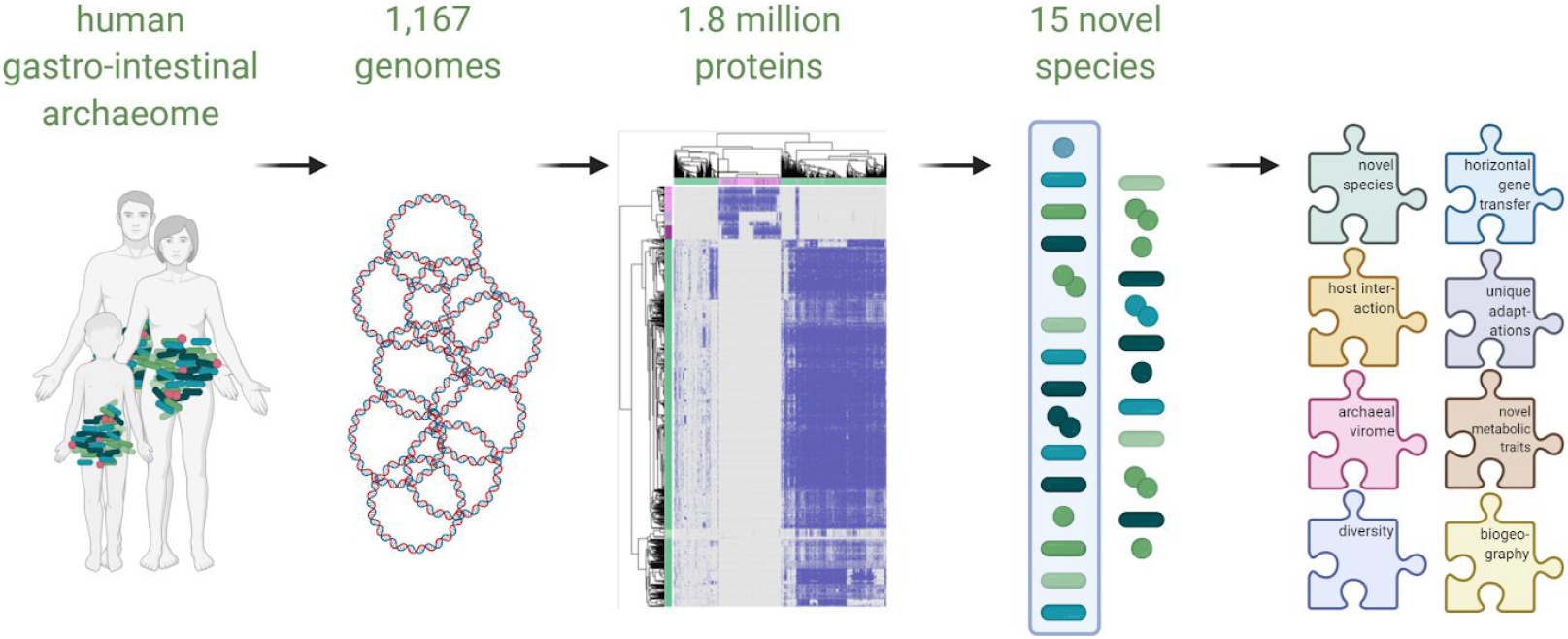

**HIGHLIGHTS:**

- The human gut archaeome analysis reveals a previously unseen active diversity
- The most abundant methanogen, *Methanobrevibacter smithii*, splits into two species
- Archaeal protein catalogue can predict geography, demographics and health aspects
- Host-associated and environmental archaea show distinct genomic & functional traits

## INTRODUCTION

The human microbiome is increasingly recognized as a key player in human health (Duvallet et al. 2017). While most research has focused on the bacterial and bacteriophages component, and to some extent unicellular eukaryotes (including fungi) and their viruses, the Archaea have been largely overlooked, mainly due to methodological reasons (Moissl-Eichinger et al. 2018; Mahnert et al. 2018; Bang and Schmitz 2018; Pausan et al. 2019; Borrel et al. 2020).

Recent analyses of metagenome-assembled genomes (MAGs) have revealed a huge unexplored human microbiome diversity, increasing the number of genome-represented bacterial species to over 2.000 (Nayfach et al. 2019; Pasolli et al. 2019; Almeida et al. 2019; 2020). Although archaea were reported in these datasets (Pasolli et al. 2019; Almeida et al. 2020), their diversity and functional potential remains largely unexplored.

Archaea are prokaryotes with distinct biology compared to bacteria and have been known to be present in the human gut for several decades (Borrel et al. 2020). Due to their regulatory function at the end of the microbial food chain, they are considered metabolic keystone species in the gastrointestinal microbial network. Some archaea also carry specific adaptive traits for survival and colonization of the human gut environment, such as bile salt hydrolases (Gaci et al. 2014), or adhesin-like proteins (Hansen et al. 2011; Borrel et al. 2017). Archaea can remove deleterious bacterial metabolites such as TMA (Brugère et al. 2014; Borrel et al. 2017) and can induce host immune responses (Bang et al. 2014; Vierbuchen et al. 2017). However, their role in human disease remains unclear (Mahnert et al. 2018; Borrel et al. 2020).

Here, we present a comprehensive analysis of a set of 1,167 archaeal genomes derived from the human gastrointestinal tract. We identified a set of genomes representing known and unknown archaeal strains (n=98) and species (n=27) from the human gastrointestinal tract. These genomes span methanogenic and halophilic archaea with a remarkably high number of as-yet undescribed representatives of the human microbiome (52 representing novel strains, and 15 novel species). In total, we identified 1.8 million coded proteins, of which ~50% remain un-annotated (0.98 million hypothetical proteins). This protein catalogue correlated with socio-demographic characteristics of the human carriers, such as age and lifestyle. The extensive resource also revealed novel taxonomic insights, such as the obvious split of *Methanobrevibacter smithii* into two separate species. Moreover, we could show that the archaeal microbiome component is actively replicating within the human gastrointestinal tract and is characterized by a specific genomic and functional adaptation towards the human host. The human gut archaeome carries a complex virome, which revealed an unexpected host flexibility for some viruses. Despite the newly available information, the archaeome still remains undersampled, indicating an even larger archaeal diversity to be expected.

This genome set will now serve as a foundation for future comparative genomics, species-directed cultivation attempts, and for further elucidating the role of the human archaeome within the microbiome and in human health and disease.

## RESULTS

### Over 1,000 unique archaeal genomes recovered from human gastrointestinal samples

To explore the diversity of archaea in human gastrointestinal samples, we compiled publicly available genomes from four recent metagenome-assembled genomes (MAGs) collections (Nayfach et al. 2019; Pasolli et al. 2019; Almeida et al. 2019; 2020) together with genomes from cultured archaea available in the NCBI (Kitts et al. 2016), PATRIC (Wattam et al. 2017) and IMG (Chen et al. 2019) repositories. To complete the dataset, we included genomes of *Candidatus* Methanomethylophilus alvus (Borrel et al. 2012), *Candidatus* Methanomassiliicoccus intestinalis (Borrel et al. 2013), of the human isolate *Methanobrevibacter arboriphilus* ANOR1 (Khelaifia et al. 2014b), and of Methanomassiliicoccales Mx02, Mx03, Mx06, and *Candidatus* M. intestinalis (Borrel et al. 2017). Henceforth, unless otherwise stated, “genomes” refers to genome sequences from both isolates and MAGs. For a detailed overview on the methodology please refer to Supplementary Fig 1.

To ensure high quality of the dataset, a threshold of >50% genome completeness and <5% contamination was used, following previously published methodology (Almeida et al. 2020). This procedure yielded 1,167 non-redundant archaeal genomes (Mash distance threshold of 0.001, 99.9% nucleotide identity; Ondov et al. 2016; Supplementary Table 1) which were further sub-grouped into a “strain”-list (<99% ANI similarity; Supplementary Table 1; 98 genomes), and a “species”-list (<95% ANI similarity; Supplementary Table 1; 27 genomes) (Fig. 1A). For this, the best quality genome (genome completeness, minimal contamination, strain heterogeneity and assembly continuity based on the N50 value) from each cluster was selected as representative, or whenever an isolate was available it was preferred and used for further analysis. More details on genome length, number of contigs, N50, G/C content, number of tRNAs and completeness are provided in Fig. 1, B-C.

**Fig. 1:**
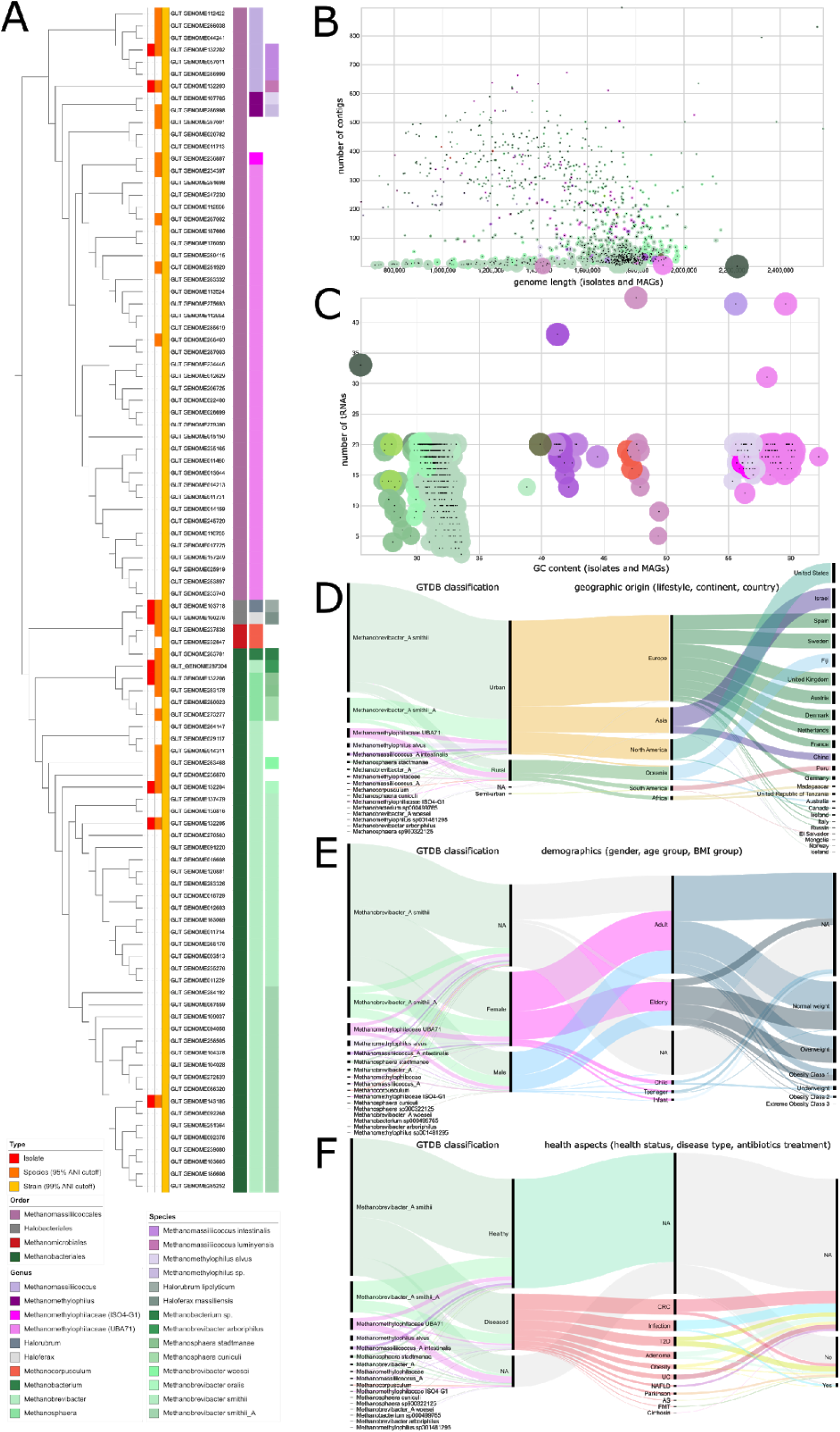
**1,167 archaeal genomes from the human gastrointestinal tract reveal taxonomic expansion of the archaeome and correlation with cohort metadata.** Note: *Methanobrevibacter smithii*_A is later referred to as *Cand*. M. intestini. **A)** ANI-based tree of genomes clustered at 99% similarity (“strains”). Isolates, species and strains are highlighted next to the tree topology, taxonomic affiliation is displayed next to the genome names. **B,C)** Genome characteristics and numerical metadata (genome length, number of contigs, N50, G/C content, number of tRNAs and completeness) are displayed as scatterplots (colored radius refers to N50 in B and completeness [%] in C). **D-F**) Categorical metadata is grouped in three alluvial diagrams referring to geographic origin (lifestyle, continent, country), demographics (gender, age group, BMI group) and health aspects (health status, disease type, antibiotics treatment) (NA = no data available).

The 1,167 genomes span a wide taxonomic diversity, and include members of the Methanobacteriales (87.15%), Methanomassiliicoccales (12.43%), Methanomicrobiales (0.26%) and Halobacteriales (0.17%; Fig. 1; Fig. 2, B-C). Most genomes were taxonomically affiliated with the known genus *Methanobrevibacter* (996 genomes; 85%), in agreement with earlier reports (Borrel et al. 2020). Other genomes were affiliated to *Methanomethylophilus* (38; 3.3%), *Methanomassiliicoccus* (29; 2.5%), *Methanosphaera* (20; 1.7%), and *Methanocorpusculum* (3; 0.3%). *Methanobacterium, Haloferax* and *Halorubrum* were only represented by one genome each. Of the 1,167 genomes, 78 (6.7%) could not be assigned to any previously described genus and 101 genomes (8.7%) did not match any known species. Those genomes are mostly Methanomassiliicoccales representatives.

**Fig. 2:**
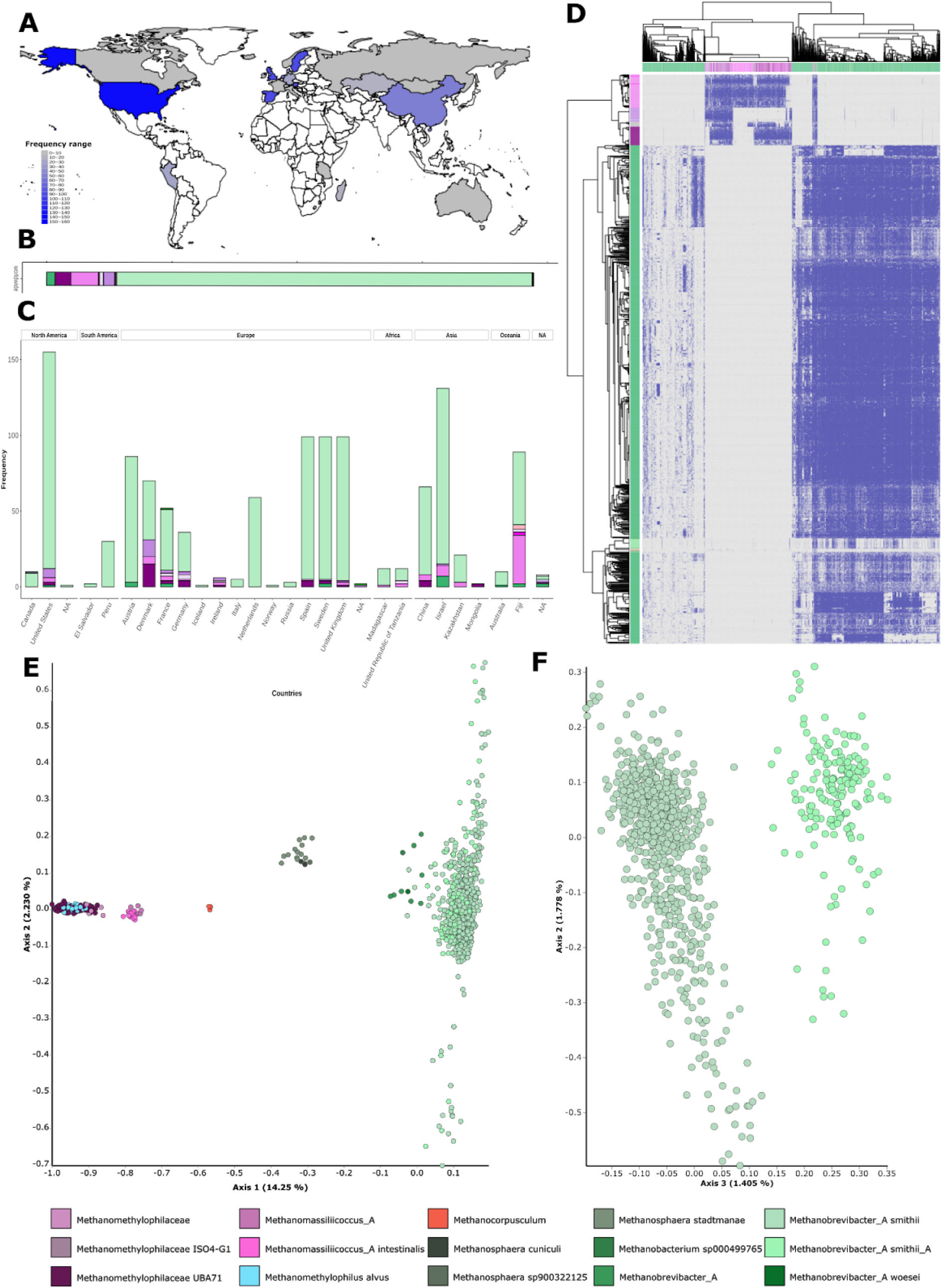
**Archaeal genomes from the human gut microbiome distribution and the corresponding unified protein catalogue.** Note: *Methanobrevibacter smithii*_A is later referred to as *Cand*. M. intestini. **A)** Worldwide distribution of the archaeal genomes, colored according to abundance (white indicates data not available). Barplot representing **B)** the overall proportion of taxa and **C)** the number per country, of the archaeal genomes isolated from the human gut. **D)** Unified human archaeal protein catalogue based on the inter-species pan-genome of all 1,167 archaeal genomes. Heatmap depicts the presence of 3,050 proteins (found in more than 50 genomes, columns) across the 1,167 archaeal genomes (rows). **E)** The taxonomic distinction of Methanomassiliicoccales, Halobacteriales and Methanobacteriales based on the protein profile (Fig. 2D), displayed in a PCoA plot based on Bray-Curtis distances at a depth of 623 archaeal proteins. The PCoA showed five distinct clusters referring to Methanomethylophilaceae, *Methanomassiliicoccus*, *Methanocorpusculum*, *Methanosphaera* and *Methanobacteriaceae*. Notably, the clade of *Methanobacteriaceae* was subdivided into *Methanobacterium* and a heterogeneous cluster of *Methanobrevibacter* (**F**).

### Archaeal protein profile correlates with geographic and demographic parameters

The entire data set (Supplementary Table 1) covers samples from rural and urban populations of 24 countries (five continents; Fig. 2, A-C) and includes more than 800 genomes from infants, children, teenagers, adults and elderly subjects spanning all BMI (Body Mass Index) groups (underweight up to extreme obesity class 3; details on available metadata are given in Star*Methods). Three-hundred subjects showed a diseased phenotype (e.g. colorectal cancer, type 2 diabetes, infection), and received antibiotic treatment. However, most genomes (3rd quartile, 75% of all values) were obtained from healthy female adults of normal weight, living in urban areas of Europe (Fig. 1, D-F).

In total, 1.8 million proteins were identified, 54% of which lacked any functional annotation. A protein catalogue of all 1,167 archaeal genomes was generated by clustering the genes predicted across all genomes (>50% amino acid identity and >80% coverage) (Supplementary Table 2). 3,050 proteins (thereof 58% hypothetical proteins) were found to be shared among more than 50 genomes in our dataset, mirroring the taxonomic distance of the two most abundant orders, Methanomassiliicoccales and Methanobacteriales (Fig. 2D). To overcome biases introduced by potential residual MAGs contamination issues, we focused our analyses on patterns observed in two or more genomes, unless stated otherwise. Additionally, we explored protein diversity patterns and their functional characterization among isolated genomes to corroborate those observed in MAGs (Supplementary Fig. 4).

Using the protein catalogue information, the predictive potential of these metadata categories was investigated. We found a high predictive potential of the unified archaeal protein matrix composition according to genome characteristics (completeness: R = 0.97, p = 1.2*10^-131^, GC content: R = 0.99, p = 7.3*10^-224^), taxonomic classification (baseline accuracy 100%) (Fig. 2, E-F), or certain demographics such as subject age (R = 0.7, p = 2.8*10^-19^, age groups baseline accuracy 73.2%), lifestyle (baseline accuracy 93.3%) or treatment with antibiotics (baseline accuracy 95.7%). To a lesser extent, other metadata categories such as the subjects health status (baseline accuracy 81.0%), their sex (baseline accuracy 73.2%), the BMI (R = 0.4, p = 2.9*10^-5^), the geographic origin (continent: baseline accuracy 78.7%, country: baseline accuracy 62.3%) or individual disease types (baseline accuracy 67.2%) were useful to describe the composition of the unified archaeal protein matrix (Supplementary Fig. 2). The created protein catalogue can now serve as basis for future functional characterizations of the archaeome, and to identify proteins potentially involved in human health and disease.

### The genetic diversity of the gut archaeome remains undersampled

To understand the level of sampling completeness of the human archaeome, we computed the genetic diversity across the 27 representative archaeal genome species, and within those genera that had over 10 genome representatives (details given in Supplementary Table 1; Fig. 3). We excluded singletons to avoid observations from potential contaminants of a MAG. We found that more genomes are needed to have a saturated overview of the genetic diversity of the genera *Methanobrevibacter* (995 genomes, α=0.63), and of the Methanomethylophilaceae UBA71 clade (66 genomes, α=0.95) (indicated by an α <1 which means each added genome contributed new genes). However, additional species of the genera *Methanosphaera* (29 genomes, α=2), *Methanomassiliicoccus* (27 genomes, α=1.07), and *Methanomethylophilus* (38 genomes, α=1.3) would not contribute new genes (for more details please refer to Star*Methods) (Tettelin et al. 2008). Observations were further confirmed by analysis at the family level excluding singletons (Supplementary Fig. 4).

**Fig. 3:**
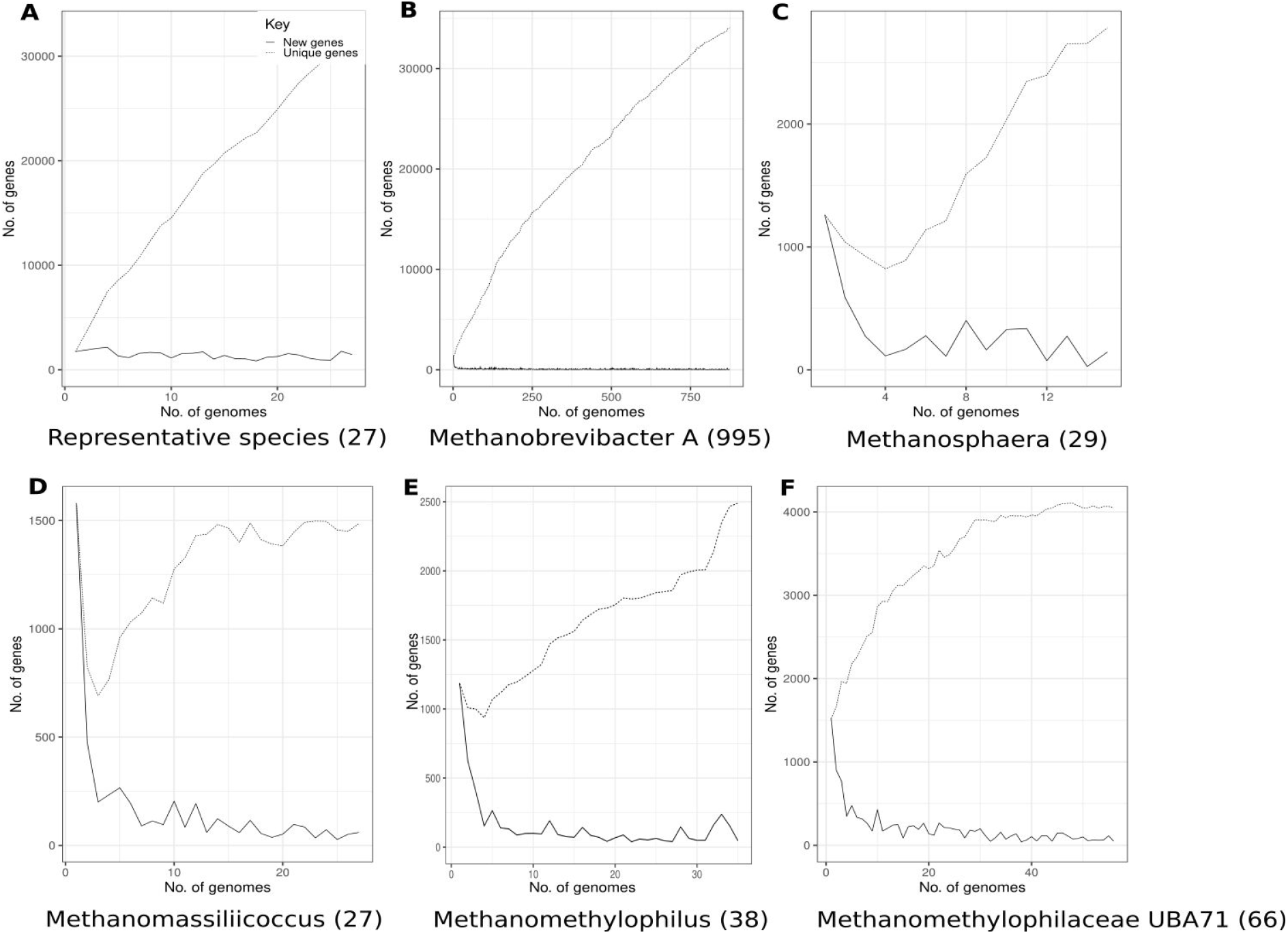
**Pan-core-genome analysis of the representative MAGs species** and for the genera that harbored over 10 genomes. The number of gene clusters for unique genes (dashed line) and new genes (solid line) results from the successive addition of genomes. The curve shows mean values of multiple iterations (> 100) where genomes were randomly added and the pan- and core-genome was computed sequentially after each addition. **A)**27 representative MAGs species, **B)** *Methanobrevibacter*_A, **C)** *Methanosphaera*, **D)** *Methanomassiliicoccus*, **E)** *Methanomethylophilus*, and **F)** Methanomethylophilacea UBA71.

Nonetheless, excluding singletons meant excluding pangenes of species which are less abundant per genus. Therefore, in analysis that included singletons, a different picture was obtained. Under these conditions, we found that more species are needed to have a saturated overview of the genetic diversity of the genera *Methanosphaera* (α=0.64), of *Methanomassilicoccus* (α=0.61), of *Methanomethylophilus* (α=0.49).

Due to the nature of MAGs, it is important to isolate more genomes of the species that are found scarcely in the dataset in order to assess the true genetic diversity of a genus in the GIT.

### The dataset reveals novel members of the human gastrointestinal archaeome, particularly in Asian and Oceanian populations

We obtained 20 genomes affiliated with *Methanosphaera*, including three genomes from isolates (Miller and Wolin 1985; Hansen et al. 2011). Taxonomically, human-associated *Methanosphaera* genomes were affiliated to three distinct species-level clades (Supplementary Fig. 5). Among those, *M. stadtmanae* was the most commonly retrieved, with 17 genomes (14 MAGs). Two MAGs (ANI 98.5%) clustered within *M. cuniculi*, and were retrieved from Asian healthy subjects living in an urban environment. The *M. cuniculi* type strain was originally isolated from the intestinal tract of a rabbit (Biavati, Vasta, and Ferry 1988) and has not been reported thus far in human hosts. One additional MAG belonging to the genus of *Methanosphaera* (Fig. 1), was binned from a gut metagenome of a diseased (colorectal cancer) European male (BMI 21, age: 64, urban environment). This genome clustered together with RUG761, a genome recovered from cattle intestines (Stewart et al. 2018); ANI 99.0%; Supplementary Fig. 5).

The dataset of human-associated Methanomassiliicoccales consisted of 145 genomes corresponding to 12 candidate species and one isolate (*M. luminyensis*) (Fig. 1, 2). The genomes were distributed into two families, the majority of them belonging to “host-associated” Methanomethylophilaceae (116 genomes), the other to Methanomassiliicoccaceae (“free-living clade”; 29 genomes). Five of the candidate species corresponded to genomes that were previously found in human samples, comprising 81% of the Methanomassiliicoccales from the current study. These included Methanomassiliicoccales Mx-06 sp. (Borrel et al. 2017; 44 genomes), *Methanomethylophilus alvus* (Borrel et al. 2012; 37 genomes) and *Methanomassiliicoccus intestinalis* (Borrel et al. 2013; 20 genomes), being the most prevalent Methanomassiliicoccales representative in human populations (Borrel et al. 2017). Mx-06 representatives were mostly present in young adults (32 years old, n = 34) from rural areas (80%; n = 40) in Oceania, Asia and Africa (65%, 13% and 7% respectively, n = 43). Together with its high prevalence (80%) in a population of 7-48 years old uncontacted Amerindians (Clemente et al. 2015; Borrel et al. 2017), it appears that this species is strongly linked with non-westernized populations. The young age of people having this species contrasts with previously reported positive correlation between age and methanogen prevalence. We found that several representatives of this species have the genetic potential to metabolize trimethylamine (TMA), a bacterial metabolite involved in trimethylaminuria and suspected in cardiometabolic, cardiovascular and renal diseases; they also have functional traits associated with adaptation to the human gut such as bile-salt hydrolase. This species is part of a well-supported clade that is separated from other Methanomethylophilaceae genus (*Methanomethylophilus*, *Methanogranum* and *Methanoplasma*) and belongs to the candidate genus “UBA71” following GTDB classification (Supplementary Table 1). We thus suggest that it represents a new genus and propose the name of *Candidatus* Methanoprimatia hominis for representatives of Mx-06 (representative MAG: GUT_GENOME268463). In addition to the species previously identified through MAGs or culture approaches, we identified six novel Methanomassiliicoccales species, represented by 24 MAGs. One of those gathers 12 MAGs and was more often found among Asian people. We propose to name it “*Ca.* Methanoprimatia macfarlanii” (representative MAG: GUT_GENOME 251929).

In addition, a number of archaeal taxa not yet described to be constituents of the human gastrointestinal tract were recovered from the MAG dataset. This included one MAG affiliated to *Halorubrum lipolyticum* (Halobacteriaceae, GUT_GENOME103718, Fig. 1), which showed 100% ANI similarity to the type strain originally isolated from a Chinese lake (Cui et al. 2006). Together with isolate *Haloferax massiliensis*, which was retrieved from human feces in 2018 (Khelaifia et al. 2018), they represent the only two genomes affiliated to halophilic archaea available from the human GIT. This observation is in contrast to the previously reported high prevalence of Haloarchaea of Asian cohorts (Kim et al. 2020). None of the two genomes revealed obvious adaptations towards the human gut ecosystem such as bile salt hydrolases and may therefore be transient residents.

Two genomes affiliated to Methanocorpusculaceae (Methanomicrobiales) (Fig. 1; ANI 98.6%), representing two strains and one species, were retrieved from subjects from the Fiji islands (Fig. 2). Closest relatives were *M. parvum* (Strain XII, type strain; ANI 70.32%) and *M. bavaricum* (DSM4179; 70.25%), which were originally isolated from digester and wastewater environments, respectively.

Several novel genomes were also detected within the *Methanobrevibacter* clade, including the cluster containing GUT_GENOME014311 (Fig. 1, Fig. 4A). This cluster was represented by seven genomes (95% ANI cut-off), clustering into two distinctive strains and one species. All seven genomes were originally binned from samples obtained from Asia (China) and Oceania (Fiji) only; genomes were obtained from healthy and diseased, rural and urban subjects. The closest relatives identified were *Methanobrevibacter woesei* GCA 003111605T (71.54% ANI) and *Methanobrevibacter smithii* ATCC 35061 NC 009515T. Additional *Methanobrevibacter* genomes were recovered with larger distance to known isolates, such as GUT_GENOME236870 and GUT_GENOME237437 (99.67% ANI; both isolated from a Fiji cohort; Fig. 1), with a genome similarity of 84.13% to *M. gottschalkii* DSM11977 (ANI), or GUT_GENOME237054 (also from a Fiji cohort), showing a similarity of 78.11% to *M. gottschalkii* (ANI). An overview on host association, geography, genome size and taxonomic association of known *Methanobrevibacter* species and genomes is given in Fig. 4, B-G). It shall be mentioned that no MAG corresponding to *Methanobrevibacter oralis* was identified in our dataset, although this species from the human oral microbiome has been at least recovered once from human stool (Khelaifia et al. 2014a).

**Fig. 4:**
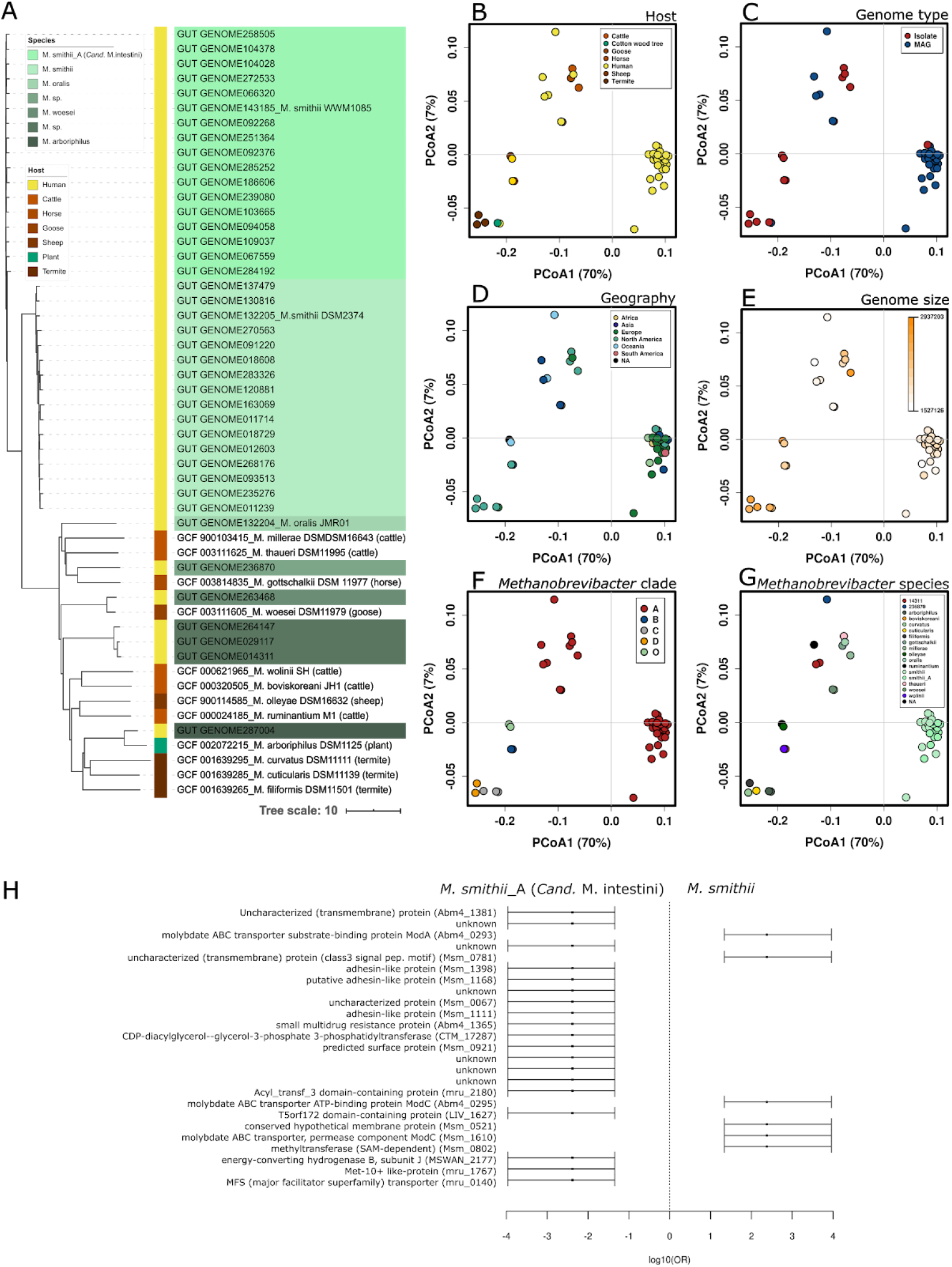
Characteristics of the *Methanobrevibacter* genomes. **A)** Phylogenetic tree of the *Methanobrevibacter* clade based on ANI distance. Twelve representative genomes from other sources than humans were included for comparison (Supplementary Table 3). Genomes (strain-level) from the human gastrointestinal tract are highlighted in green colors (taxon label). The bar on the left displays the origin: human (yellow bar), animals (shades of red) and plant (green). **B-G)** PcoA plots of protein profiles according to host **(B)**, genome source type **(C)**, geography **(D)**, genome size (**E**), *Methanobrevibacter* clade according to GTDB **(F)** and species **(G)**. **H)** Forest plot showing the outcome of the Wilcoxon rank test comparison of genomes from *M. smithii*_A (*Cand*. M. 16S rRNAintestini) vs. *M. smithii* (only TOP 25 proteins are shown; FDR<0.000005); bar displays the odds ratio (OR) (Supplementary Table 4).

Signatures of the genus *Methanobacterium* have been frequently detected in amplicon-based archaeome studies using small intestine biopsies or samples from oral cavities, but have neither been isolated nor detected by genome-centric analyses (Faveri et al. 2011; Matarazzo et al. 2011; Koskinen et al. 2017). The presence of *Methanobacterium* species in the human GIT have been confirmed in our studies, as one Methanobacterium MAG (GUT_GENOME283701) was obtained from a European male (age 65; colorectal cancer), with a genome size of 1.9 Mbp (98.13% completeness; 0% contamination). Based on ANI values, the closest described relative was *Methanobacterium formicicum* DSM3637 (87.75%), a well-known, formate consuming methanogen in ruminants (Chellapandi et al. 2018). Formate consuming capacity was as well found for the *Methanobacterium* MAG identified in our study.

Undersampled populations obviously bear an unknown archaeal wealth which requires future dedicated studies.

### The *Methanobrevibacter smithii* clade splits into two separate speciesh

The *Methanobrevibacter smithii* clade was the largest taxonomic clade detected in our study (Fig. 2). Based on ANI similarity values, as well as information derived from the protein catalogue, the *Methanobrevibacter smithii* group was represented by two species-level clades (tentatively named “smithii” and “smithii_A” according to GTDB classification; Fig. 2F, Fig. 4A; Supplementary Table 1 also Pasolli et al. 2019). *M. smithii*_A was represented in our entire dataset 185 times (16% of entire dataset), whereas *M. smithii* was detected 797 times (68%), together representing 84% of all genomes in our dataset (Fig. 2, A-C). Notably, the two *M. smithii* groups differed significantly with respect to their protein profile (PERMANOVA, p=0.001; Fig. 2F) and genome size (p<0.001, Wilcoxon rank sum test, two-sided, genome size corrected by completeness), with median genome sizes of 1.7 Mbp for *M. smithii* and 1.8 Mbp for *M. smithii*_A (Supplementary Table 4; genome sizes for isolates: 1.7 Mbp (reference genome *M. smithii*) and 1.9 Mbp (isolate WWM1085, see below), respectively).

All *M. smithii* strains carried the *modA* gene, which was not detected in any of the “smithii_A” genomes (Supplementary Table 4). This gene is involved in molybdate transport, and responsible for substrate binding (Self et al. 2001). In addition, amongst the top 25 discriminative proteins (Fig. 4H), the molybdate ABC transporter permease component, as well as the molybdate ABC transporter ATP-binding protein was identified in 94% of all *M. smithii* genomes, but in none of the *M. smithii*_A genomes. This indicates a different pathway for molybdate acquisition in the *M. smithii*_A clade. The *M. smithii*_A genomes were further characterized by additional unique membrane/cell-wall associated proteins, such as adhesin-like proteins, surface proteins, and a number of uncharacterized membrane proteins (Fig. 4H).

Based on the extent of discriminative features, and an average nucleotide identity of 93.95% between the two representative genomes of *M. smithii* and *M. smithii*_A, we propose to rename the smithii_A clade, represented by isolate WWM1085 (GUT_GENOME143185, Jennings et al. 2017), into *Candidatus* Methanobrevibacter intestini to further emphasize the presence of two predominant, distinctive *Methanobrevibacter* clades in the human intestinal tract. *Cand*. M. intestini and *M. smithii* cannot be distinguished on 16S rRNA gene sequence, which might be the reason for missing this separation in two distinctive clades before. However, analysis of the *mcrA* gene revealed a consistent difference between the two clades, with an average of 2.15 % difference in amino acid sequence (1.82-2.22%; Supplementary Material 1).

In future studies it will be important to further investigate the distribution and the potential interplay of *M. smithii* and *Cand.* M. intestini, in order to understand their biological role(s) in the human body.

### The human archaeome is actively replicating

In order to address the question whether the human gastrointestinal archaea are actively replicating in their host environment, we performed iRep analysis (see Star*Methods). Based on iRep analysis of 26 genomes with fully available metagenomic data resources, 10-50% of the archaeal population was found to be actively replicating at the time point of sampling with a mean replication rate of 1.4+/−0.1. Replication rates differed between *M. smithii, M. smithii_A* (*Cand.* M. intestini)*, Methanomassiliicoccus*, and Methanomethylophilaceae UBA71 clade. The highest replication rate was observed for *Methanobrevibacter*, the lowest was observed for the *Methanomethylophilaceae* representatives; this finding, however, was not statistically significant (Kruskal-Wallis test on all groups: p = 0.4, pairwise Wilcoxon rank sum test on selected taxa: mean FDR corrected p value = 0.615). Also, iRep values did not reveal any significant correlations with geographic origin, demographics or health aspects (Supplementary Fig. 3). In case of unfiltered iRep values from 69 genomes, the mean replication rate increased to 1.9+/−0.9. According to this data set the highest replication rate was observed again for *Methanobrevibacter*, but here the lowest was observed for *Methanomassiliicoccus*. Interestingly, both subsets revealed higher replication rates of *Methanobrevibacter smithii_A* (*Cand*. M. intestini) than for *Methanobrevibacter smithii* (Supplementary Fig. 3).

### The human archaeome carries a complex, previously unseen virome

The virome associated with gut archaea is largely unknown. We identified up to 94 viral populations in our genome datasets by using CheckV (Nayfach, Camargo, et al. 2020). This number is the result of clustering 45 high-quality (>90% completeness) and 130 medium quality (50-90% completeness) archaeal proviruses at 95% identity and 80% coverage. The selected cut-off is repetitively used and is shown to be consistent with viral species definition (Duhaime and Sullivan 2012; Brum et al. 2015; Duhaime et al. 2017; Bobay and Ochman 2018; Roux et al. 2019; Gregory et al. 2016; 2020) (Fig. 5; Supplementary Table 5). Of the identified high-quality (HQ) proviruses, 91 viral species representatives were found to be specific for *Methanobrevibacter*, one for *Methanomassiliicoccus*, one for *Methanosphaera*, and one for Methanomethylophilaceae UBA71.

**Fig. 5:**
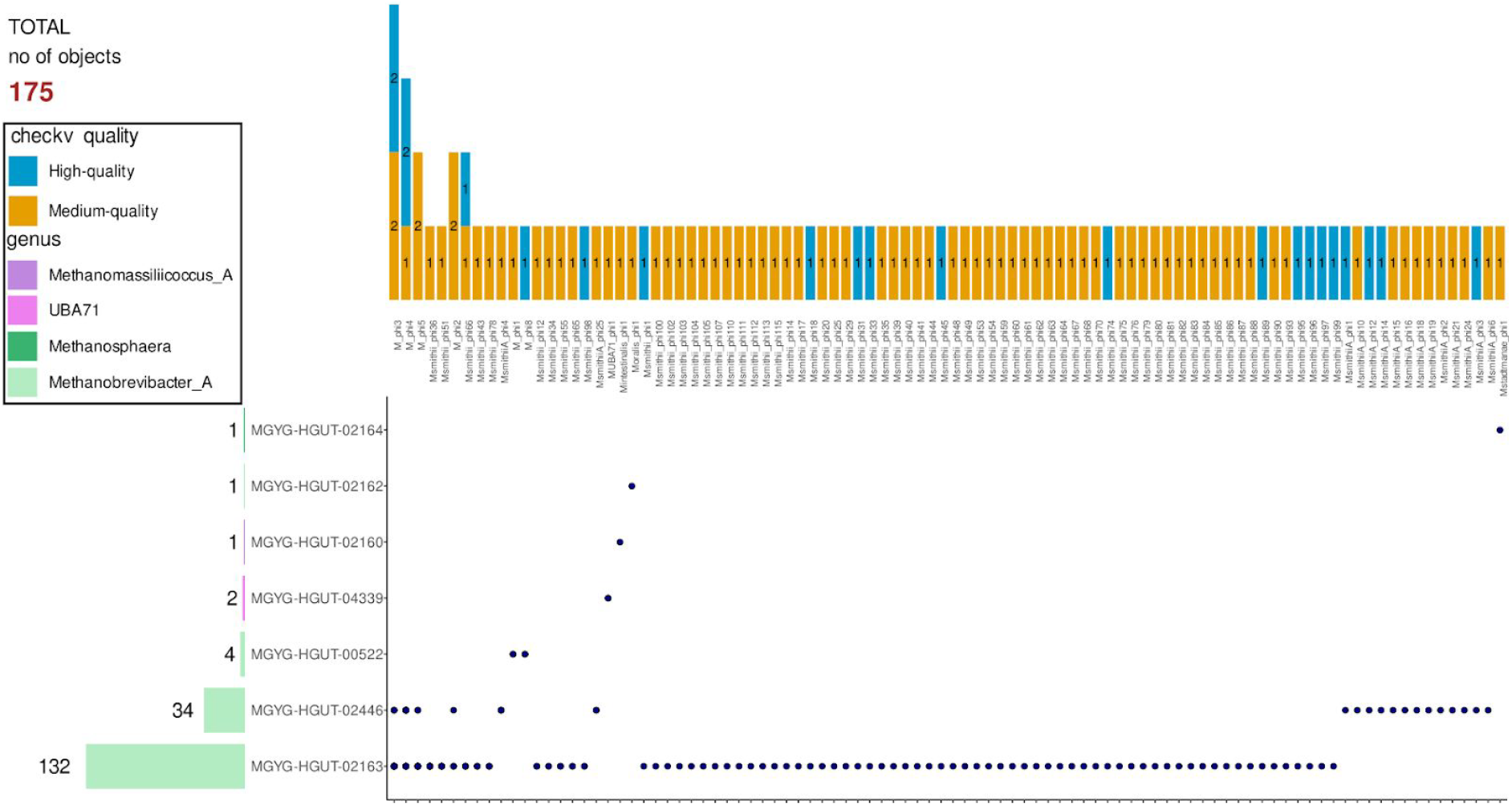
UpSet plot showing presence of viral species representatives color coded by the assigned quality (yellow for high-quality, and blue for medium-quality) in genome species where a member of the viral cluster was identified. Viral clusters representative were named as follows: if the cluster is of viruses infecting hosts of the same genus the name is Mgenus_phi_Number and if the cluster is of viruses infecting hosts of different genera the name is M_phi_Number.

Notably, the identified proviruses seem to infect i) the same host regardless of geographical location which suggests that those viruses can be widespread, and ii) hosts of different species (specifically, prophages infecting both *M. smithii, Cand.* M. intestini, Supplementary Table 5) which suggests a broad host range.

While archaeal viruses in extreme environments have been discovered in the early 1970s (Torsvik and Dundas 1974; 1980), little is known about non-extremophilic viruses in the highly abundant mesophilic environments and only a few non-extremophilic archaeal viruses have been isolated so far (review and overview papers of archaeal viruses Snyder et al. 2015; Prangishvili et al. 2006; 2017). Only a handful of archaeal viruses have been described to infect the **same archaea genera** widespread in both the environment and the human host. In more detail, we found no similarities between the identified proviruses and the lytic phages Drs3 infecting *Methanobacterium formicicum* (Wolf et al. 2019), six viruses infecting various *Halorubrum* species (Pietilä et al. 2012), the provirus Msmi-Pro1 infecting *Methanobrevibacter smithii* ATCC 35061 (Krupovič, Forterre, and Bamford 2010) and the provirus φmru infecting *Methanobrevibacter ruminantium* (Ouwerkerk, Gilbert, and Klieve 2011). However, to the best of our knowledge, no viruses/proviruses have been identified in the past infecting *Methanomassiliicoccales* and *Methanobacteriales* members of the human gut.

We additionally explored the novelty of these proviruses by comparing them with the latest comprehensive human Gut Virome Database (GVD) (Gregory et al. 2020), and the Viral Refseq Database using the network-based viral classification tool vConTACT2 (Bin Jang et al. 2019). No clustering resulted between the high- and medium-quality identified viruses and both databases. Due to the lack of similar archaeal viral genomes in the reference databases, the classification and further characterization of discovered archaeal viruses through metagenomic approaches remains challenging.

Taken together, these results reveal that archaeal viruses probably have a currently underestimated diversity and likely ecological importance in the human gut microbiome. Consequently, the current and ongoing identification of archaeal viral species provide unique opportunities for future in-depth functional characterizations, taxonomic classification, estimation of the prevalence of each viral species, isolation and follow-up studies of virus-archaea dynamics.

### Human-associated archaea exhibit a lower proportion of horizontal gene transfer than animal-associated archaea

The adaptation of archaea to the human gut environment may have been favored by specific acquisition of genes from the resident bacterial community providing new functions. To assess this possibility, we compared the retrieved *Methanosphaera* and *Methanobrevibacter* genomes (strain list, only isolates and MAGs with 0% contamination) with isolates and genomes derived from animal sources (Supplementary Table 6; Fig. 6).

**Fig. 6:**
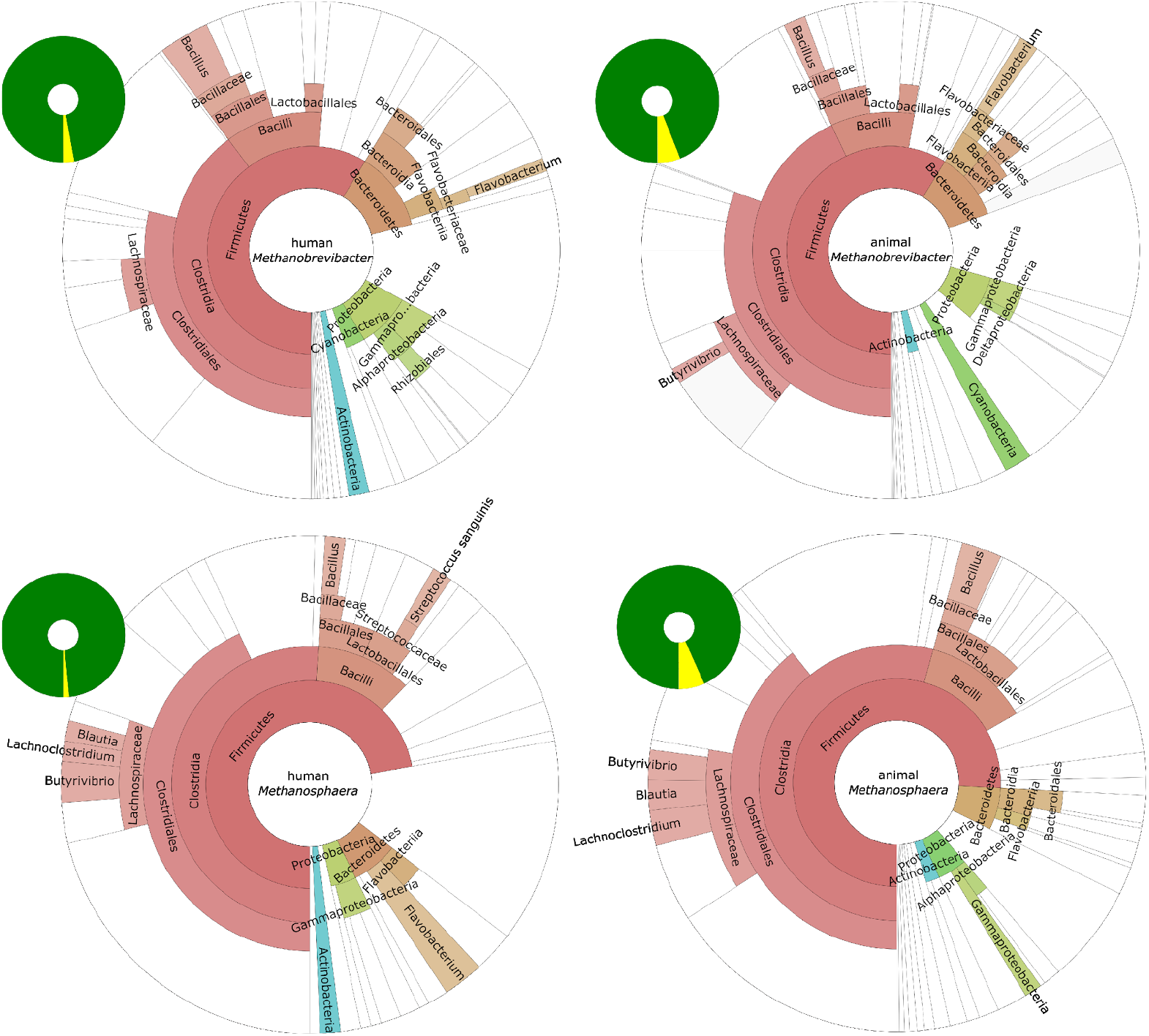
**Contribution of bacterial-annotated genes** in human- (left) and animal- (right) associated *Methanobrevibacter* and *Methanosphaera* species: proportion (small circles, yellow refers to proportion of bacterial annotation) and potential bacterial origin (taxa as displayed in the large circles). Unclassified taxa are whitened out. Only MAGs with 0% contamination and of high quality (taken from “strain list”) and genomes from isolates were analyzed.

Human-associated methanogens revealed a significantly lower proportion of non-archaeal genes, irrespective of whether we considered isolates only or both isolates and MAGs. Human-associated *Methanobrevibacter* species carried on average approx. 2.84% genes annotated as of non-archaeal origin, which was significantly lower than the proportion of non-archaeal genes in animal-associated *Methanobrevibacter* species (6.09%; Mann-Whitney U test, p=0.00308; genomes from isolates only: 6.36%). In particular *Methanobrevibacter* smithii/smithii_A (*Cand.* M. intestini) representatives revealed a very low contribution of non-archaeal genes (2.11%; genomes from isolates only: 1.8%). Human-associated *Methanosphaera* species carried on average a proportion of 1.45% of genes of bacterial annotation (genomes from isolates only: 0.68%). Animal-associated *Methanosphaera*, however, contained a significantly higher proportion of bacterial genes (6.74%; p=0.00452, Mann-Whitney U test; genomes from animal isolates only: 5.31%).

An extraordinary high contribution of bacterial genes were observed for Methanomassiliicoccales, revealing a proportion of archaeal-annotated genes of 74.96% only (mean of 9 MAGs (0% contamination, high quality genomes; strain list) and 2 isolates). The proportion ranged from 65.72-96.88% in the MAGs, the genomes from isolates revealed an archaeal-gene annotation of 67.35 and 69.65%. This might reflect however an incomplete annotation of the Methanomassiliicoccales genomes based on the low number of available genomes and isolates.

In all cases, the largest contribution was observed from Firmicutes (Clostridia and bacilli; example bile salt hydrolases see below) with a lower contribution of Bacteroidetes, and Proteobacteria. Those clades represent the most abundant bacterial microbiome components, increasing the probability for HGT towards the archaeome (Lurie-Weinberger et al. 2012) (Fig. 6; Supplementary Material 3).

Our results indicate that adaptation towards the human host is not necessarily reflected by a higher proportion of genes derived from the human gastrointestinal bacteriome.

### Host-associated archaea diverge from environmental relatives

We reasoned that host-associated archaea diverge taxonomically and functionally due to the characteristics of their individual host environments.

In 16S rRNA gene-based analyses (details given in Star*Methods), we found that members of genera *Methanobrevibacter* and *Methanosphaera*, as well as *Cand*. Methanomethylophilus belonged almost exclusively to taxa from host-associated (animal, human, plant) sources, whereas *Methanocorpusculum*, Nitrososphaeria and Haloferaceae were more related to environmental strains (Fig. 7A). ANI-based analyses of the families Methanobacteriaceae, Methanocorpusculaceae, Methanomethylophilaceae and Methanomassiliicoccaceae revealed an overall clear separation between the MAGs of different origins (Fig. 7, B-E). More specifically, for Methanomassiliicoccaceae (Fig. 7C), we observed that the two gut genomes classified as *M. luminyensis* (GUT_GENOME132203 and GUT_GENOME140888) cluster within an environmental archaeal MAGs clade whose members are taxonomically classified as uncultured *Methanomassiliicoccus*; however, both human-derived genomes share a higher ANI value compared to the genomes of the sister clade. Concerning Methanobacteriaceae (Fig. 7E), GCA_002509095.1 (Methanobacteriaceae archaeon UBA254) and GUT_GENOME283178 (*Methanosphaera* sp900322125) shared an ANI value of 96.09% while every other pairwise comparison between human and environmental archaeal MAG had an ANI value less than 88.09%.

**Fig. 7:**
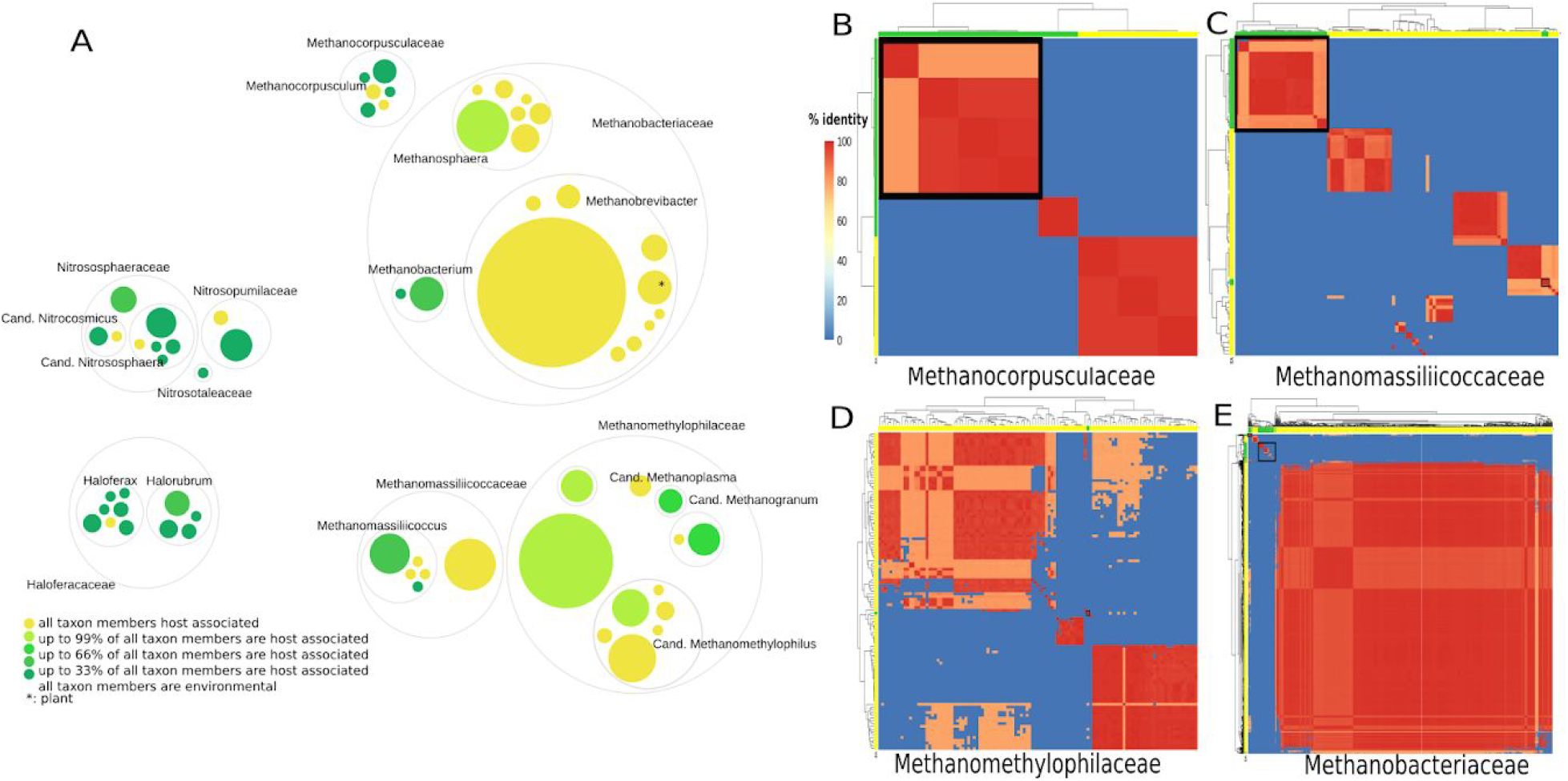
Comparison of host-associated and environmental relatives. **A)** Circle packing plot, displaying the environmental or host-associated nature of specific taxa. Analysis performed on 16S rRNA gene level. Number of sequences analyzed per taxon are indicated by circle size; colors indicate the proportion of host-associated signatures (see legend). The largest contribution was observed from *Methanobrevibacter smithii* sequences (n= 192). Note, that the yellow color (“host-associated”) also includes human, animal and plant (*; only *M. arboriphilus*)-associated taxa. **B-C-D-E)** Average nucleotide identity (ANI) percent identity heatmap visualization. ANI analysis based on MinHash sequence mapping was performed using fastANI and visualized using the pheatmap library in R. ANI values represented range from 75–80% ANI colored in light orange, 80-90% ANI colored in darker orange, and over 95% ANI in red. Heatmap for genomes assigned to the taxonomic family of **B)** Methanocorpusculaceae, **C)** Methanomassiliicoccaceae, **D)** Methanomethylophilaceae, and **E)** Methanobacteriaceae. Genomes isolated from the human gut microbiome (labeling on the x and y-axes in yellow) can be separated from the genomes isolated from the environment (labeling on the x and y-axes in green; Supplementary Table 7). Environmental archaeal genome clustering is marked with a black square.

Based on taxonomic classification resolved at the species level, most species identified in the human gut are indeed host associated (reviewed in Bang and Schmitz 2018; Moissl-Eichinger et al. 2018; Borrel et al. 2020). More specifically, based on the information on their respective biomes, the archaeal genomes of this study can be classified into three groups: i) exclusively found in the human gut, ii) host (human, animal, plant)-associated and iii) widespread in the environment, with the first two groups representing the majority (reviewed in Bang and Schmitz 2018; Moissl-Eichinger et al. 2018; Borrel et al. 2020). Following this classification and based on the current availability of genomes and metadata, *H. massiliensis*, *M. oralis*, *M. smithii*, *M. smithii_A* (now: *Cand*. M. intestini), *M. stadtmanae*, *M. intestinalis* and *M. alvus* can be considered to be affiliated to group (i). Species belonging to group (ii) include *M. woesei*, and *M. cuniculi*. Species of group (iii) are represented by *H. lipolyticum* (Cui et al. 2006), *M. arboriphilus* (Zeikus and Henning 1975; Khelaifia et al. 2014b) and *M. luminyensis* (Dridi et al. 2012; Borrel et al. 2017), which are widespread in various kinds of environments.

It should be noted that for certain clades only a limited number of genomes is currently available, and thus their host/environmental tropism remains to be precisely determined (e.g. Methanomethylophilus sp001481295, *Methanosphaera* sp900322125, *Methanobacterium* sp000499765, *H. massiliensis*, *Methanomassiliicoccus)*. Notably, in both types of analyses, the genus *Methanobrevibacter* was overwhelmingly affiliated with the human and other hosts. This is a unique trait amongst all analyzed archaeal genera (Fig. 7A).

### Metabolic and functional interaction of the archaeome with the gut environment

We analyzed specific features that could indicate the advanced interaction of the human-associated GIT archaea with their gut environment, i.e. with host and non-archaeal microbiome components (Fig. 8).

**Fig. 8:**
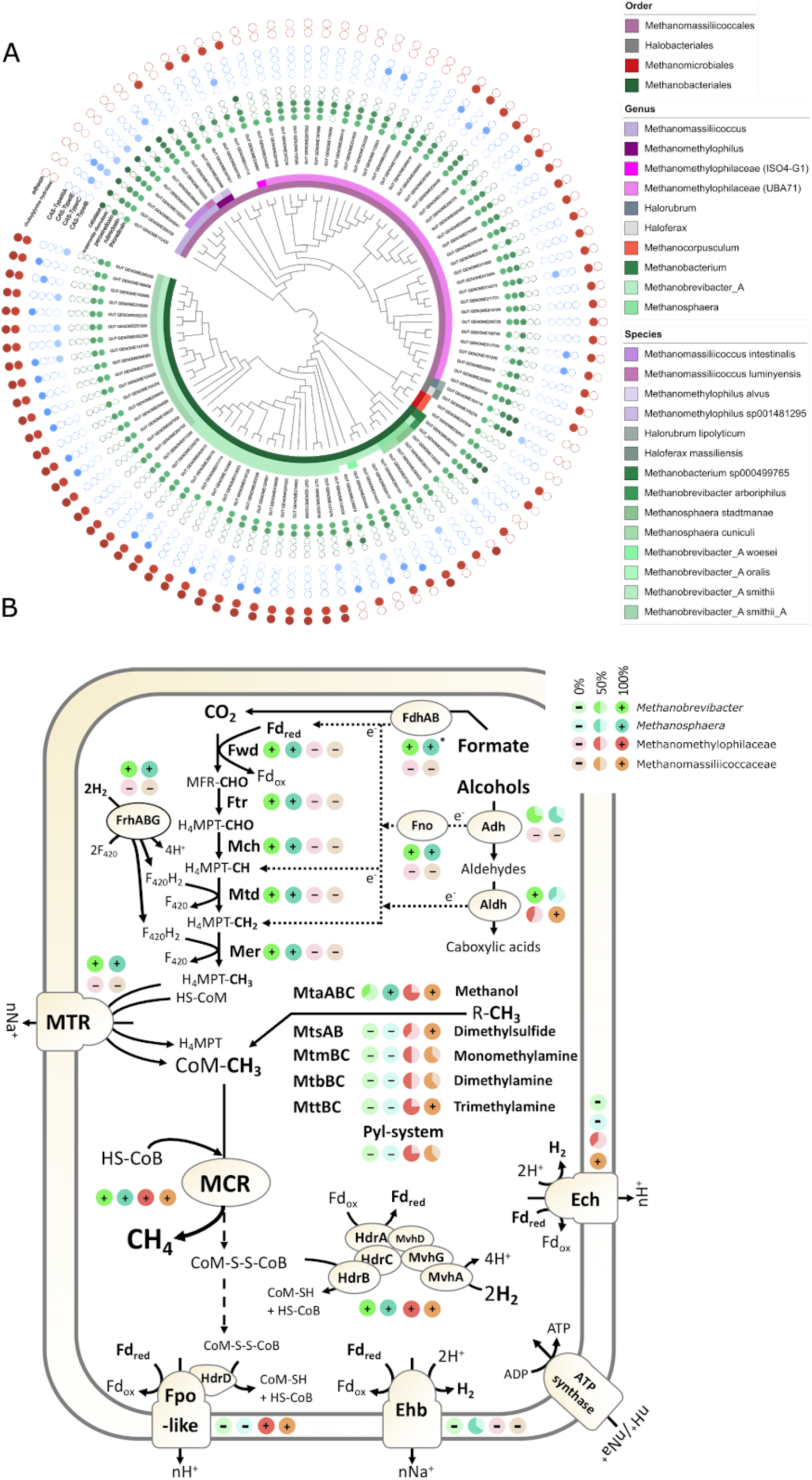
**Adaptations of human gut archaea with respect to ecosystem and metabolism.** Note: M. smithii_A corresponds to *Cand*. M. intestini. **A)** Overview of the absence and presence of genes involved in host-interaction (red), oxygen resistance (green) and detected CAS systems (blue). Genomes (strain-list) were analyzed using MaGe Microscope and genes were counted as present when automatic annotation was positive (“putative” annotation was counted as positive). **B)** Methanogenic pathways in 23 human gut-associated Methanobacteriales and Methanomassiliicoccales species. The proportion of species with a given protein or protein complex is indicated by pie charts for *Methanobrevibacter* (n=7), *Methanosphaera* (n=3), Methanomethylophilaceae (n=8) and *Methanomassiliicoccus* (n = 5). For clarity, the nature of the electron transporter and some intermediate steps in the electron transfers are not displayed for formate and alcohol utilization. R-CH3 corresponds to methanol, dimethylsulfide, monomethylamine, dimethylamine or trimethylamine. Alcohol could be ethanol or secondary alcohols. The absence of certain enzymes may be due to the fact that the MAGs are incomplete.

Loss of genes involved in dealing with oxidative stress are considered as a trait of host-association, as environmental strains have to face non-permanently, strict anaerobic conditions, while this is not the case for strains inhabiting the GIT. We therefore specifically analyzed the presence of genes associated with oxygen resistance (catalase, superoxide dismutase, peroxiredoxin, rubredoxin and thioredoxin; Lyu and Lu 2018). Catalase was detected in some Methanomassiliicoccales (mainly *Methanomassiliicoccus* representatives) and Haloarchaea, and in *Methanobrevibacter arboriphilus* and *Methanobacterium*. The presence of a superoxide dismutase was rarely detected, namely in *Haloferax* and *Halorburum* members. None of the *Methanobrevibacter* representatives, except *M. arboriphilus*, carried the peroxiredoxin gene. In contrast, thioredoxin and rubredoxin were detected in the majority of the genomes.

Additional functions of interest are adhesins and bile salt hydrolases (i.e. choloylglycine hydrolase, CGH). A number of adhesins was specifically annotated in the genomes of *Methanobrevibacter* and *Methanosphaera* species (Fig. 8A). CGH homologues were detected in 11 out of 28 of the archaeal species, including the five most prevalent ones (*M. smithii*, *Cand.* M. intestini, *M. stadtmanae*, *M. alvus* and *Cand.* M. hominis). CGH genes were not detected in any of the Methanomassiliicoccus genomes and in the Haloferaceae, indicating their importance for specialization toward the human gut. It should be noted that the CGH genes detected in Methanomassilicoccales, Methanomicrobiales and Methanobacteriales formed separate clusters within Firmicutes bile salt hydrolases gene tree (Supplementary Fig. 6), indicating their potential acquisition from different events of HGTs.

Additional adaptations were observed at the metabolism level. Apart from key components of methanogenesis, MCR and Hdr/Mvh complexes, the main gut methanogens (Methanobacteriales and Methanomassiliicoccales) possess very distinct methanogenesis pathways (Fig. 8B; Supplementary Table 8). For example, differently to all Methanomassiliicoccales, all human gut *Methanobrevibacter* have the genetic potential for formate and H2/CO2 utilisation. However, both Methanobacteriales and Methanomassiliicoccales representatives carry the *mtaABC* genes, providing the genetic potential to use methanol. In total, 83% of the MAGs have these genes, suggesting that methanol utilization might provide a selective advantage in the human gut. However, the conditions under which *Methanobrevibacter* use methanol and whether this compound is a methanogenesis substrate or enters an anabolic pathway remain unknown.

The two dominant *Methanobrevibacter* species display the genetic potential to use alcohols (likely secondary alcohols and ethanol) as electron donors for methanogenesis. One of the *Methanosphaera* species may also have the genetic capacity to reduce methanol with ethanol for methanogenesis as described earlier (Hoedt et al. 2016), but this species was only encountered once in our analyses, and *Methanosphaera stadtmanae*, the main *Methanosphaera* species in human, cannot perform this pathway.

The majority (11/13) of the human-gut associated Methanomassiliicoccales species code for the MttBC methyltransferase and corrinoid protein needed for methanogenesis from TMA. This capacity would allow them to decrease the concentrations of this molecule produced by gut microbiota and involved in cardiovascular diseases (Brugère et al. 2014; Borrel et al. 2017). The presence of the *mttBC* genes was detected in a larger proportion of the Methanomassiliicoccales MAGs originating from Europe and North America (~60%) with respect to Africa and Asia (~40%) or Oceania (17%) (Supplementary Fig. 7). These variations may reflect different TMA-production capacity by bacteria in the microbiota across these populations and diet habits. One of the two Methanomethylophilaceae species lacking TMA-utilization capacity (*Cand*. Methanoprimatia macfarlanii) also lacks MtbBC and MtmBC methyltransferases and corrinoid proteins for dimethylamine and monomethylamine utilization, respectively. However, several strains of this species have the genes to encode Pyrrolysine synthesis (*pylSBCD*), a proteinogenic (UAG codon-encoded) amino-acid almost only present these methylamine methyltransferases (Srinivasan, James, and Krzycki 2002; Brugère et al. 2018). The absence of detection of the methylamine methyltransferases in these MAGs, including MttBC for TMA utilization, is thus likely due to their incompleteness. The other species lacking methylamine methyltransferase, corresponding to Methanomassiliicoccales Mx02 sp. (Borrel et al. 2017), also lacks any other genes known to be involved in methyl-compound utilization or in any alternative methanogenesis pathways (Supplementary Table 8). The absence of these methanogenesis genes in all the MAGs of Methanomassiliicoccales Mx02 sp. and in previously obtained related MAGs, support assumptions (Borrel et al. 2017; Söllinger et al. 2016) on the presence of novel methanogenesis pathways likely based on unknown methyltransferases, or another metabolic route in the Methanomassiliicoccales. We propose the name of *Candidatus* Methanarchanum hacksteinii Mx02 for this potential new genus and species (representative MAG: GUT_GENOME287001). Our observations indicate that members of the human archaeome possess a variety of specific adaptations towards this specific environment, many of which are probably not yet discovered.

## DISCUSSION

This work adds novel information on the biology of the human gastrointestinal archaeome, by characterizing an extended collection of 1,167 non-redundant archaeal genomes. This assessment of the genome collection and the catalogue of 1.8 million putative proteins can now serve as a reference base for extensive investigation of the hitherto unexplored gastrointestinal archaeome.

The comprehensive analysis of the globally distributed gut archaeome offers a unique resource to generate hypotheses for future studies. This includes aspects on the archaeal physiology, the interaction with the bacterial microbiome and the virome, the comparison of animal- and human-host association, and the archaeal cross-talk with the human host. Moreover, the provided resource can serve as an important database for targeted cultivation of members of the archaeome, as well as their unique virome.

One of the main remaining questions is whether archaeal pathogens do exist, since a per se pathogenic capacity of archaea has never been identified (Borrel et al. 2020). Therefore, our resource is an important basis for further exploring potential pathogenic traits, such as virulence factors.

As our insights are largely predicted from genomic information, additional work is urgently required to address the archaeome functional capacity using expression and cultivation-based assays. Furthermore, our dataset is based on publicly available samples, which were originally processed for the analysis of the bacterial component of the microbiome (Almeida et al. 2019; Nayfach et al. 2019; Pasolli et al. 2019; Nayfach, Roux, et al. 2020; Almeida et al. 2020). Therefore, archaeal species that require specialized methods (Mahnert et al. 2018) for cell lysis and DNA extraction may be missing from our current collection. Moreover, as several taxa in our collection are only represented by single genomes, additional conspecific strains will be needed to validate many of our genomic insights.

Future efforts should seek to extend the dataset beyond the gastrointestinal environment, to other human body sites (Koskinen et al. 2017) and hosts (Moissl-Eichinger et al. 2018). Moreover, incorporating both transcriptomics and proteomics data in the future will further reinforce the genomic predictions and improve our understanding of the regulation of archaea physiology and host adaptation.

Overall this work contributes substantially to the understanding of the microbiome as a complex multi-domain microbial network of the human gastrointestinal tract (Taffner et al. 2018; Nayfach, Roux, et al. 2020; Youngblut, Reischer, et al. 2020; Youngblut, Cuesta-Zuluaga, et al. 2020). We showed that despite the availability of more than 1,000 genomes, the human archaeome still remains undersampled. Future work aiming to expand the archaeal pangenome will enable the identification of specific archaeal functions/molecular mechanisms that may be causal in microbiome-associated human health conditions.

## ACKNOWLEDGEMENTS

CME received funding from the Austrian Science Fund FWF (P 32697, P 30796), which is highly appreciated. CME and AM gratefully acknowledge computational resources of the MedBioNode at the Medical University Graz and the Life Science Compute Cluster (LiSC) operated by the Computational Systems Biology group at the University of Vienna. We thank the Medical University Graz ZMF Galaxy Team: Core Facility Computational Bioanalytics, Medical University of Graz, funded by the Austrian Federal Ministry of Education, Science and Research, Hochschulraum-Strukturmittel 2016 grant as part of BioTechMed Graz. We gratefully acknowledge the discussion with Katharina Bick on taxonomic naming, and the scientific support provided by Zobia Hameed.

RS received partial funding from Kiel Marine science cluster at the CAU and the German ministry of education and science, BMBF (031B0851B), which is highly appreciated. CC and AW used resources of the high-memory computer nodes and computation support from the Computing Center of the CAU Kiel. AA and RF are supported by EMBL.

JFB acknowledges a grant from Hub Innovergne (“Investissements d’Avenir” 16-IDEX-0001 CAP 20–25).

GB and SG acknowledge support by the Institut Pasteur and the French National Research Agency (ANR) through grants Methevol (ANR-19-CE02-0005-01)and Archevol (ANR-16-CE02-0005-01).

## AUTHOR CONTRIBUTIONS

Conception and design: AA, RS, CME, SG, GB

Methodology and analysis: CC, AM, GB, CME, JFB

Collection, data processing and data generation: AA, RF, CC, AW, AM, GB

Data interpretation: CC, AM, GB, JFB, RS, CME

Manuscript preparation: CC, AM, GB, JFB, RS, CME

## STAR METHODS

### RESOURCE AVAILABILITY

#### · Lead Contact

Further information and requests for resources and reagents should be directed to and will be fulfilled by the Lead Contact Christine Moissl-Eichinger.

#### · Data and Code Availability

All the recovered genomes are available at [http://ftp.ebi.ac.uk/pub/databases/metagenomics/genome_sets/archaea_gut-genomes.tar.gz]. All the other considered genomes and metagenomes are publicly available in NCBI, and referenced. This study did not generate code, mentioned tools used for the data analysis were applied with default parameters unless specified otherwise.

## METHOD DETAILS

### · Dataset description

We selected archaeal genomes sampled from the human gut publicly available as part of the Unified Human Gastrointestinal Genome (UHGG) collection (Data access June 2020, https://www.ebi.ac.uk/metagenomics/genomes) (Almeida et al. 2020) and the MGnify resource (Mitchell et al. 2020). Briefly, this collection of non-redundant metagenomes-assembled genomes (MAGs) and isolates was collected from public repositories and metadata for each genome was gathered as well (see Almeida et al. 2020 for more details). Genomes were compared using Mash v2.1 (Ondov et al. 2016) and for genomes that were estimated to be identical, had a Mash distance of 0, only one was selected. In addition, we included genomes of ‘Ca. *Methanomethylophilus alvus’* (Borrel et al. 2012), ‘Ca. *Methanomassiliicoccus intestinalis’* (Borrel et al. 2013), as well as human gut-derived MAGs of Methanomassiliicoccales Mx02, Mx03, Mx06 and additional Ca. M. intestinalis’ (Borrel et al. 2017), and the human isolate *Methanobrevibacter arboriphilus* ANOR1 (Khelaifia et al. 2014b) to complete the dataset. Those genomes were assigned a genome accession number (GUT_GENOME286998 to GUT_GENOME287004), as given in Supplementary Table 1. This brought the total number of genomes used for the analysis in this paper up to 1,167 genomes.

### · Genome quality and taxonomic classification

The completeness of the non-redundant 1,167 genomes was evaluated by CheckM v1.0.11 (Parks et al. 2015) and only genomes which were more than 50% complete and had less than 5% contamination were selected (following the protocol from Almeida et al. 2020).

DRep v2.0.0 (Olm et al. 2017) was used to dereplicate the complete dataset at 95% and 99% average nucleotide identity (ANI). 95% ANI values were selected to separate between species boundaries (n=27) and a cutoff of 99% was chosen to distinguish archaeal MAGs on strain level (n=98) (Jain et al. 2018). The resulting strain- and species representatives are given in Supplementary Table 1.

All genomes were taxonomically annotated following the procedure given in (Almeida et al. 2020). The taxonomic assignment was performed using the Genome Taxonomy Database Toolkit (GTDB-Tk) version 0.3.1 (database release 04-RS89) (Chaumeil et al. 2020) and default parameters. Methodology is detailed in Supplementary Fig. 1.

### · Genome annotation and protein catalogue

Protein-coding sequences (CDS) were predicted and annotated with Prokka v1.14.5 (Seemann 2014) using the parameters ‘--kingdom Archaea’ to include non fragmented archaea curated proteins from the UniProtKB database and ‘--rfam’ to scan for ncRNAs. CDS were further characterized using eggNOG mapper v2.0.0 (Huerta-Cepas et al. 2017) and the eggNOG database v5.0 (Huerta-Cepas et al. 2019).

The protein catalogue was generated by combining all predicted CDS (total number 1,790,493) derived from the 1,167 non-redundant archaeal genomes. MMseqs2 linclust (Steinegger and Söding 2017) was used to cluster the concatenated proteins dataset using the options ‘--cov-mode 1 -c 0.8’ (minimum coverage threshold of 80% the length of the shortest sequence) and ‘--kmer-per-seq 80’. To reduce the risk of contaminants, the proteins were filtered to remove all non clustered proteins. This gave a total of 28,581 proteins clustering at 50% identity (Supplementary Table 2) visualized using the library pheatmap (Kolde 2019) in R.

In addition to the protein catalogue, the various species- and strain-subsets of the total 1,167 archaeal genomes (Supplementary Table 1) were submitted to MaGe MicroScope (Microbial Genome Annotation & Analysis Platform, Vallenet et al. 2009), for detailed analyses of genomic synteny.

### · Correlation with geographic and demographic parameters

Beside genomic information (genome length, number of contigs, N50, GC content, genome completeness, genome contamination, number of rRNAs and tRNAs), eleven metadata categories (numerical 2, categorical 9) could be considered for the data set. Information about the geographic origin was available for 1,063 genomes (91% of the data set, covered countries from maximum to minimum: United States, Israel, Spain, Sweden, Fiji, United Kingdom, Austria, Denmark, Netherlands, France, China, Peru, Germany, Madagascar, United Republic of Tanzania, Australia, Canada, Ireland, Italy, Russia, El Salvador, Iceland, Mongolia, Norway). Information about lifestyle was available for 1,054 genomes (90%, max-min: urban, rural, semi-urban), health state (healthy, diseased) for 894 genomes (77%), age group (adult, elderly, child, teenager, infant) for 825 genomes (71%), gender (female, male) for 620 genomes (53%), body mass index group (normal weight, overweight, obesity class 1, underweight, obesity class 2, extreme obesity class 3) for 505 genomes (43%), name of disease (colorectal cancer CRC, infection, type 2 diabetes T2D, Adenoma, obesity, Ulcerative colitis UC, nonalcoholic fatty liver disease NAFLD, Parkinson, Ankylosing spondylitis - arthritis AS, fecal microbiota transplant FMT, cirrhosis) for 303 genomes (26%), treatment (antibiotics) for 241 genomes (21%).

Supervised classification and regressions with RandomForest were applied to predict respective metadata categories from the unified archaeal protein catalogue with the q2-sample-classifier plugin (Bokulich et al. 2018). First subsets of each metadata category were created from the entire protein matrix and randomly split into a training and test set with the proportions 80 to. 20%. By using K-fold cross validation, the training set served as a learning model to predict class probabilities with settings for optimized feature-selection and parameter tuning. In the end, model accuracy was determined by comparing the predicted values between the training and test data sets.

### · Pan-genome analysis

Pan-genome analysis was performed using Roary version 3.11.2 (Page et al. 2015) using a minimum amino acid identity of 80% and no paralog splitting. Since Roary was not intended for comparing extremely diverse sets of genomes, it was used for pan-genome analysis for archaeal genomes of the same families or the same genus. We used Heaps’ Law (η = κ ∗ N−α) to estimate whether we have an open or a closed pan-genome (Tettelin et al. 2008), a first estimate for all predicted CDS and a second estimate excluding singletons from gene presence absence matrices. This analysis was carried out in the R package ‘micropan’ (Snipen and Liland 2015) using a default permutation value of 100, where η is the predicted number of genes for a particular number of Genomes (N), and κ (intercept parameter) and α (decay parameter) are the constants used to fit the curve after the genomes are ordered in a random way. An open pan-genome is indicated by an α <1 while a closed pan-genome is indicated by a α >1.

### · Estimation of replication rates

Replication rates were estimated with iRep (Brown et al. 2016) after mapping reads with Bowtie 2 (Langmead and Salzberg 2012) and post-processed with samtools (Li et al. 2009). Since the original raw reads were not available for each representative genome, as the remaining read sets were not made publically available, read coverage between the origin of replication and its terminus (main input for iRep) could only be mapped for 69 out of 98 genomes.

### · In-depth taxonomic and phylogenetic analyses of the various genera

ANI distances and tree matrices were calculated using the online resources of the enveomics platform (Rodriguez-R and Konstantinidis 2016), MaGe (Vallenet et al. 2009), as well as Microbial Genomes Atlas (MiGA) (Rodriguez-R et al. 2018). Phylogenetic trees, based on the ANI tree matrix, were annotated using the iTOL tool (Interactive Tree Of Life) (Letunic and Bork 2019), and processed using InkScape. For specific considerations involving additional genomes from animals, a subselection of the archaeal genomes was reanalyzed together with the additional genomes following the same settings as described for the protein catalogue procedures above (respective datasets are given in the Supplementary Table 3 and Supplementary Table 9 as described in the main text).

McrA genes were extracted via MaGe, hosting all strain level genomes. McrA genes were aligned using MegaX (Kumar et al. 2018), and a maximum likelihood tree was calculated (default settings).

### · Horizontal gene transfer analysis

The information from protein catalogue incl. eggNOG annotation was used for the assessment of the horizontal-gene transfer candidates. For this, the taxonomic “best hit” information for all annotated genes was used; only genomes revealing 0% contamination were used for this assessment. All data are provided in Supplementary Table 6. Data were visualized using Krona (Ondov, Bergman, and Phillippy 2011).

### · Detection of virulence and resistance genes

To predict potential virulence genes in all 1,167 archaeal genomes, ABRicate version 0.5 (https://github.com/tseemann/abricate) was used to profile the following databases: CARD (Jia et al. 2017), Resfinder (Zankari et al. 2012), PlasmidFinder (Carattoli et al. 2014), ARG-ANNOT (Gupta et al. 2014), EcOH (Ingle et al. 2016), MEGARes 2.0 (Doster et al. 2020) as well as NCBI AMRFinderPlus (Feldgarden et al. 2019). Since ABRicate is solely based on DNA sequences, blastX searches using DIAMOND (Buchfink, Xie, and Huson 2015) was used to complement results from ABRicate on the level of protein sequences in the virulence factor database (VFDB version 20191122) (Chen et al. 2005; Liu et al. 2019) and CARD together with the Resistance Gene Identifier (RGI) (Alcock et al. 2020).

Specific groups of proteins and genes involved in human interaction were investigated according to available annotations from MaGe (Vallenet et al. 2009) and eggNOG-mapper (Huerta-Cepas et al. 2017).

### · Viral identification, quality estimation and comparisons to viral databases

To assess the presence of prophages, and estimate completeness and quality of potential proviruses, CheckV (Nayfach, Camargo, et al. 2020) was used to scan all MAGs. To ensure we overcome possible contamination issues that can potentially result from the binning process, we selected proviruses flanked within archaeal contigs for this analysis. CheckV looks for host-virus boundaries based on difference in GC content and gene annotation in a sliding window approach. Proviruses that had a CheckV quality assignment of medium-quality (50-90% completeness), high-quality (>90% completeness) or complete were considered for further analysis. Quality assignments by CheckV are based on Minimum Information about an Uncultivated Virus (MIUViG) standards (Roux et al. 2019). The selected phages were subsequently clustered with MMseqs2 using the ‘linclust’ function with the same parameters previously specified and Mmseqs function ‘result2repseq’ to select a viral cluster representative.

ORFs of viral populations with the previously specified MIUViG quality were used as input for vConTACT2 (Bin Jang et al. 2019) including Viral RefSeq genome (version 97). VconTACT2 is used to affiliate a family or a genus rank group to viral populations and thus to determine taxonomic diversity.

A recent study was published by Gregory et al. 2020 (Gregory et al. 2020) where a human Gut Virome Database (GVD) harbors 33,242 viral populations including 0.1 % archaeal viruses resulting from 2,697 gut metagenomes from 32 studies. This dataset was used as a reference database to scan the identified viral scaffolds using MMseqs2 “easy-linclust’’ function at 50, 80, 90 and 95% identity.

In addition, we BLASTED (50% identity and 80% coverage cut off) identified proviruses to characterized proviruses and lytic phages infecting members of the same genera we find in our dataset. The proviruses Msmi-Pro1 (coordinates 1693232–173205) infecting *Methanobrevibacter smithii* ATCC 35061 (Accession Number CP000678) (Krupovič, Forterre, and Bamford 2010), and the provirus φmru infecting *Methanobrevibacter ruminantium* were used (Ouwerkerk, Gilbert, and Klieve 2011). Additional lytic phage Drs3 infecting *Methanobacterium formicicum* (Wolf et al. 2019), σM1 infecting *Methanothermobacter thermoautotrophicum* (Meile et al. 1989), and six viruses infecting various *Halorubrum* species (Pietilä et al. 2012) were used for comparison.

### · Comparison to environmental archaea

For considerations based on 16S rRNA genes, 16S rRNA genes of representative genomes were extracted using Metaxa2 (Bengtsson-Palme et al. 2015) (n=314; not all 16S rRNA genes could be recovered). This dataset was supplemented with data from amplicon sequencing studies and clone sequences from archaeal signatures from human gastrointestinal samples (dataset described in Borrel et al, 2020; n=381 in total). These sequences were aligned and classified using the SILVA rRNA database (Quast et al. 2013). More specifically, the retrieved 16S rRNA genes were subjected to the ACT tool (alignment, classification and tree service) (Carver et al. 2005), using the following parameters: Basic alignment parameters: “removed”; Search and classify, min. identity with query sequence: “0.95”, Number of neighbours per query sequence: “10”; Compute tree, Workflow: “Denovo including neighbours” and default parameters; Advanced tree computation parameters, Positional variability filter: “None”, Domain: “Archaea”. Unclassified sequences were removed from the dataset. Via SILVA SINA, 10 next neighbours were selected, and information on their isolation source was gathered through NCBI (Supplementary Material 2; Supplementary Table 7; the final dataset contained 566 sequences). Grouping was performed on genus/species level, and information on the percentage of host-associated archaea in all groups was displayed as a circle packing plot (RawGraphs online tool, https://app.rawgraphs.io/).

For genome-based analyses, a set of 623 archaeal MAGs identified from environmental and gastrointestinal samples (for example, rumen, guinea pigs and baboon faeces) was used as a reference dataset for comparison to the set of archaea isolated from the human gut microbiome (Parks et al. 2017; Almeida et al. 2020). All environmental genomes used were as well over 50% complete up to 90% complete with less than 5% contamination as well. To estimate the pairwise Average Nucleotide Identity (ANI) distance between environmental archaeal genomes dataset (Supplementary Table 7) and the archaeal genomes from the human gut microbiome, we used fastANI (Jain et al. 2018), a tool that effectively discriminated intra- and inter-species boundaries for over 90K prokaryotic genomes.

### · Metabolic interaction of the archaeome with the gastro-intestinal environment

Proteins involved in methanogenesis were searched in all genomes using custom Hidden Markov Models (HMM) profiles (threshold e-value 10^-5^) implemented in Macsyfinder (Abby et al. 2014). This allowed to determine the presence of enzymatic complexes on the basis of the presence of all or most subunits. The presence in the 26 methanogenic species was first evaluated based on the representative genome (that are the most complete/less contaminated). If a majority of the MAGs in a species have an enzyme, then this enzyme was considered as present in the species, even if absent from the best representative genome.

### · Functional interaction of the archaeome with the gastro-intestinal environment (exploring oxygen resistance, adhesins and bile salt hydrolases)

Specific functions were searched (“search by keywords”-function) in MaGe (Vallenet et al. 2009) (all strain genomes have been made publicly accessible). Presence and absence information was used for tree annotation through iTOL (Letunic and Bork 2019). The backbone tree was based on ANI similarity as described above.

#### · Tools used for data visualization

PCoAs and other graphical displays based on the unified archaeal protein matrix were calculated and visualized in Qiime2 (Bolyen et al. 2018) and Calypso (Zakrzewski et al. 2017). Venn diagrams were created with creately (https://creately.com/). Alluvial plots and circle packing plots were generated with RAWGraphs (https://app.rawgraphs.io/). Strip charts were created with Calypso. Phylogenetic trees, based on the ANI tree matrix, were annotated using the iTOL tool (Interactive Tree Of Life). All figure panels were created using InkScape.

## QUANTIFICATION AND STATISTICAL ANALYSIS

All statistical analyses were conducted by using R, Qiime2 and Calyspo. Statistical significance was determined by non-parametric tests including Spearman rank correlations, PERMANOVA, Wilcoxon rank sum tests for pairwise analysis, Mann-Whitney U tests for unpaired data and Kruskal-Wallis tests if significance had to be determined for all groups. Significance was considered at an alpha <0.05 after 999 permutations. P-values were corrected for multi hypothesis testing using the false discovery rate (FDR).

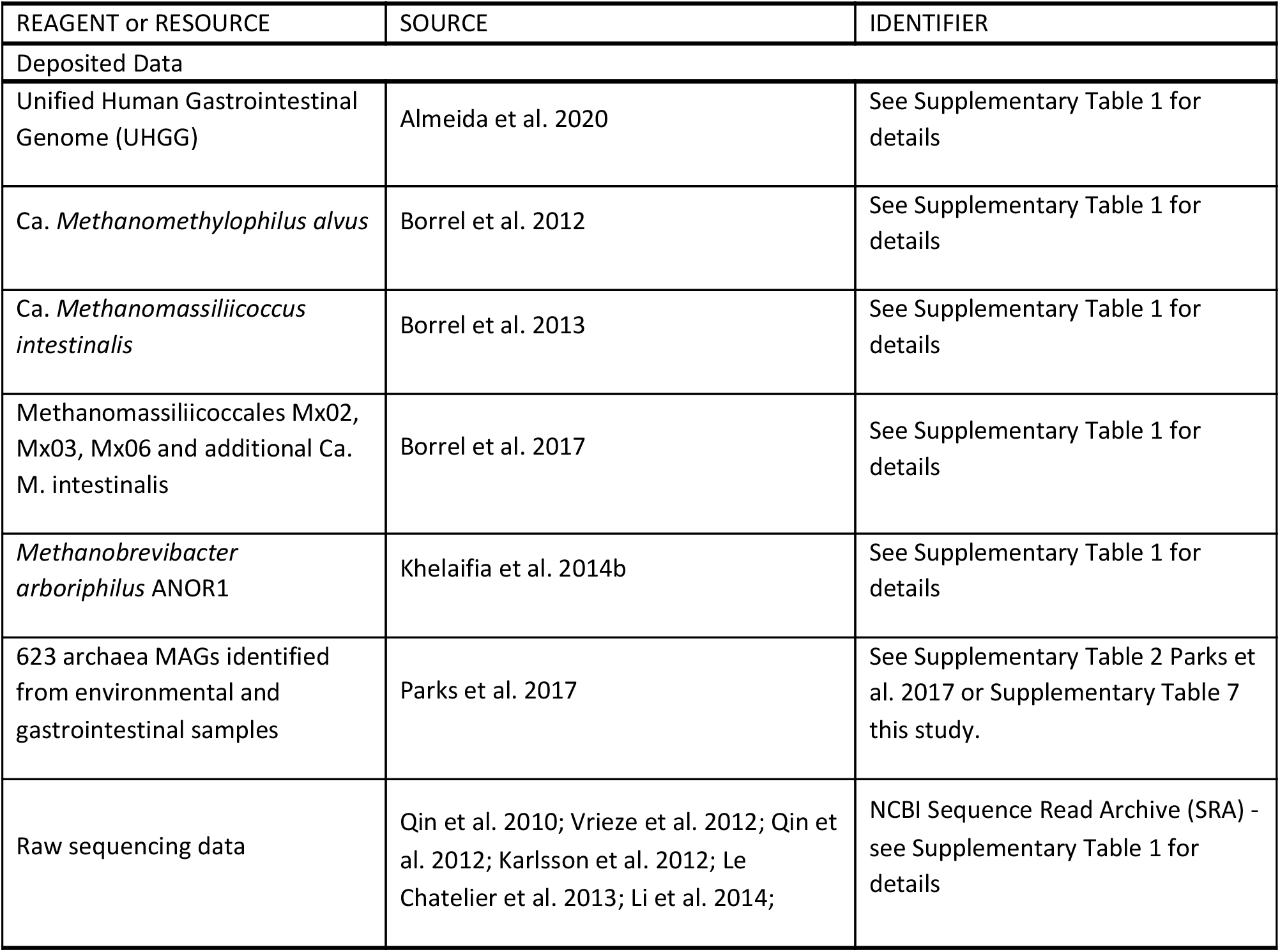
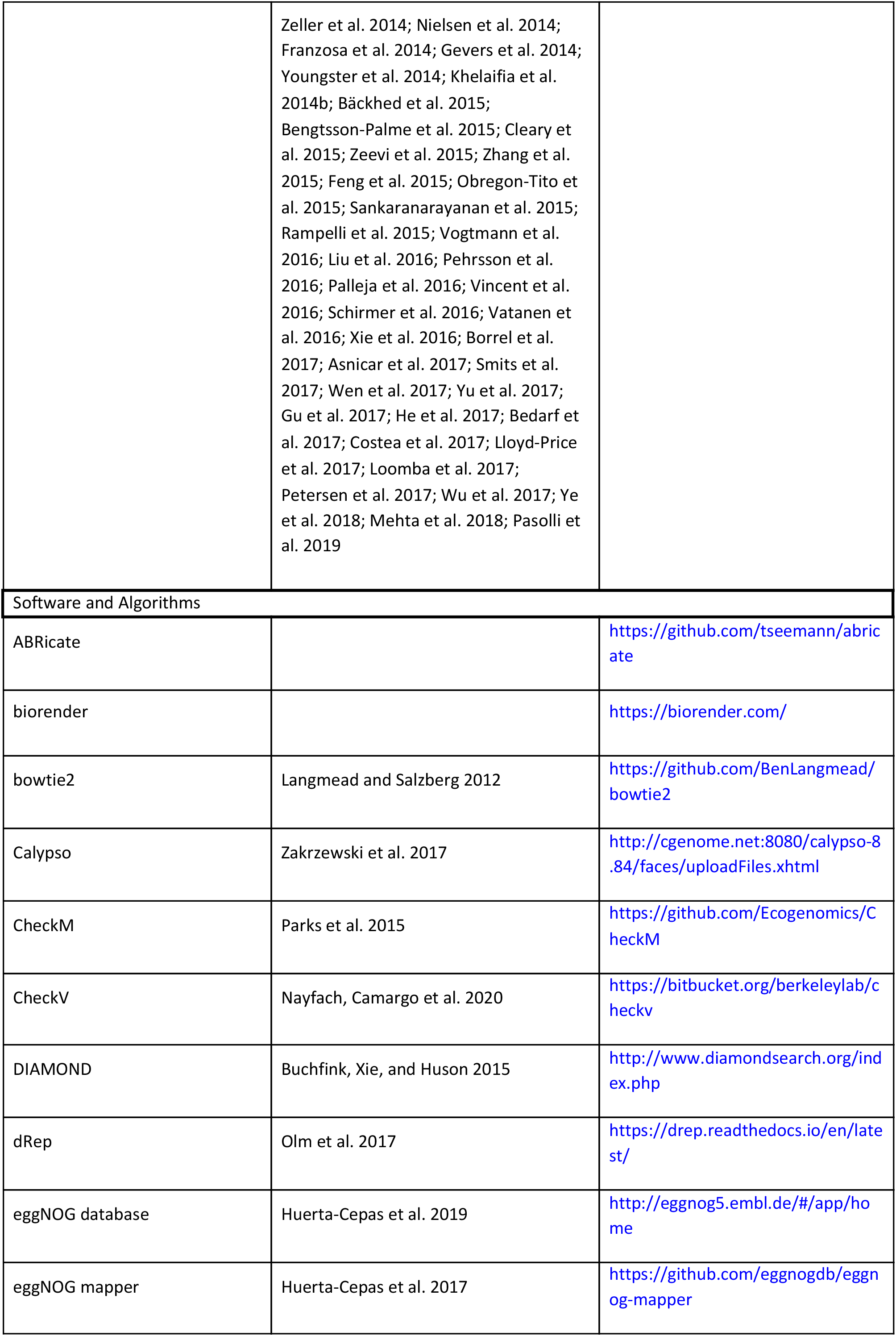
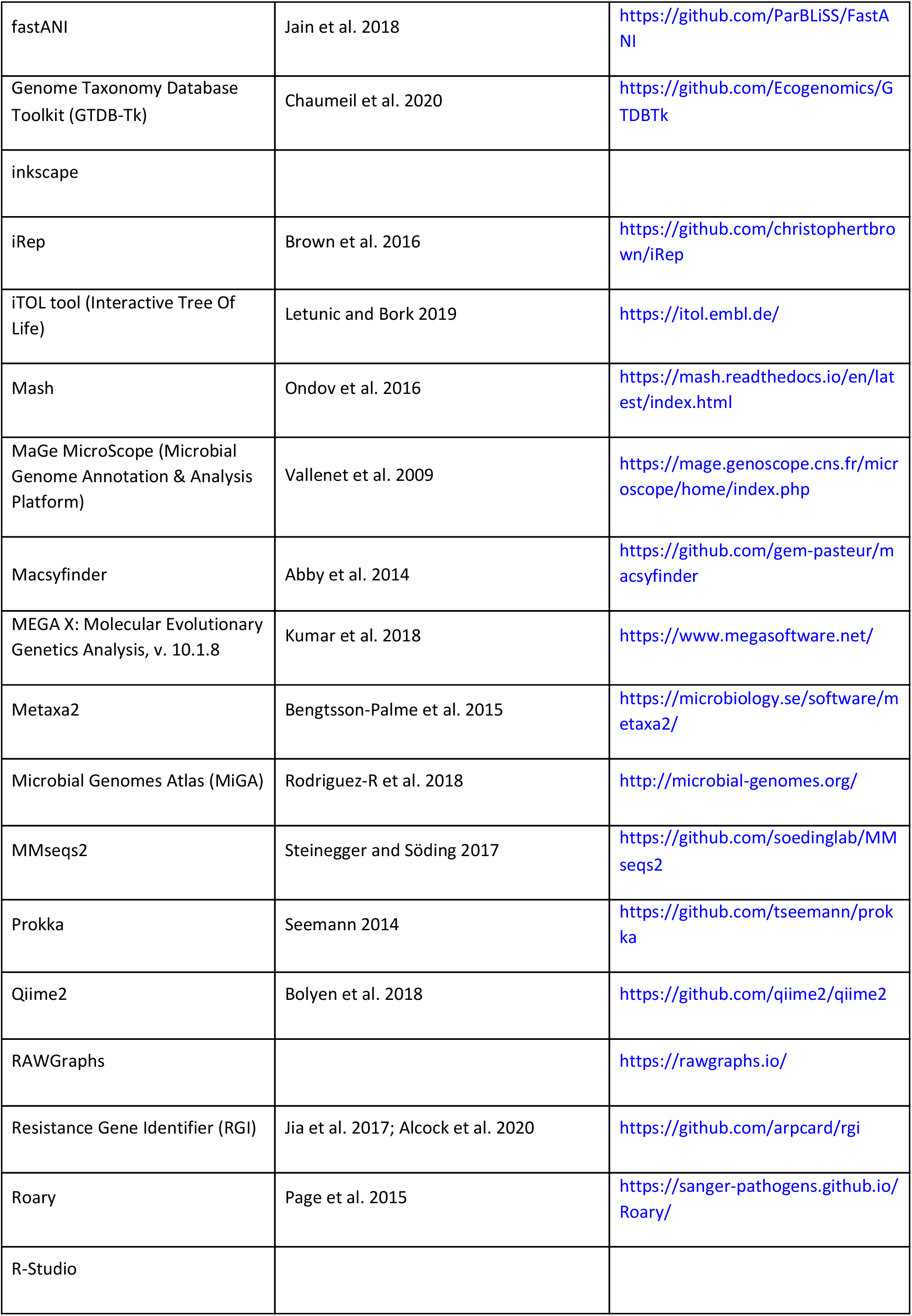
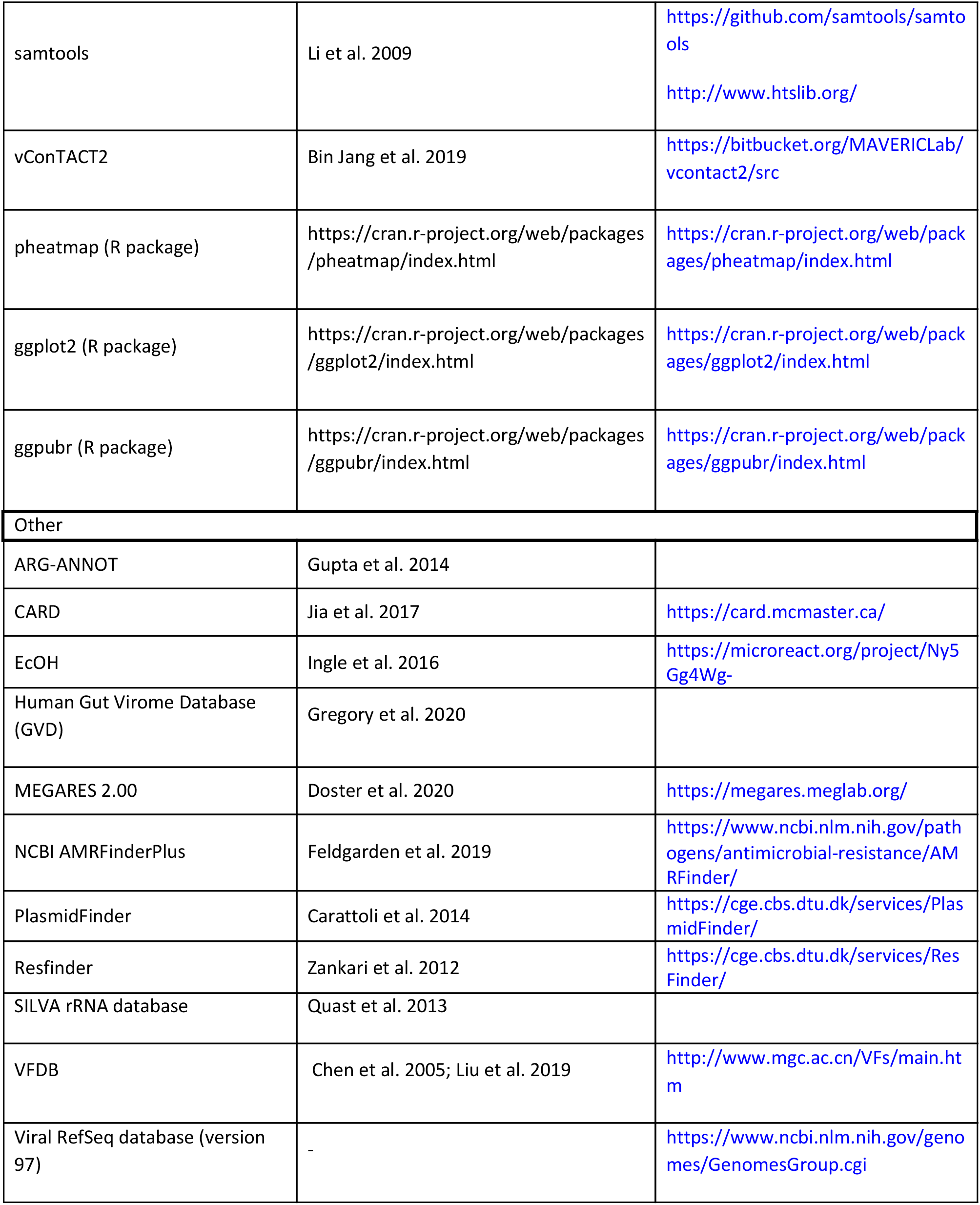
KEY RESOURCES TABLE.

## SUPPLEMENTAL INFORMATION

**Supplementary Fig. 1:**
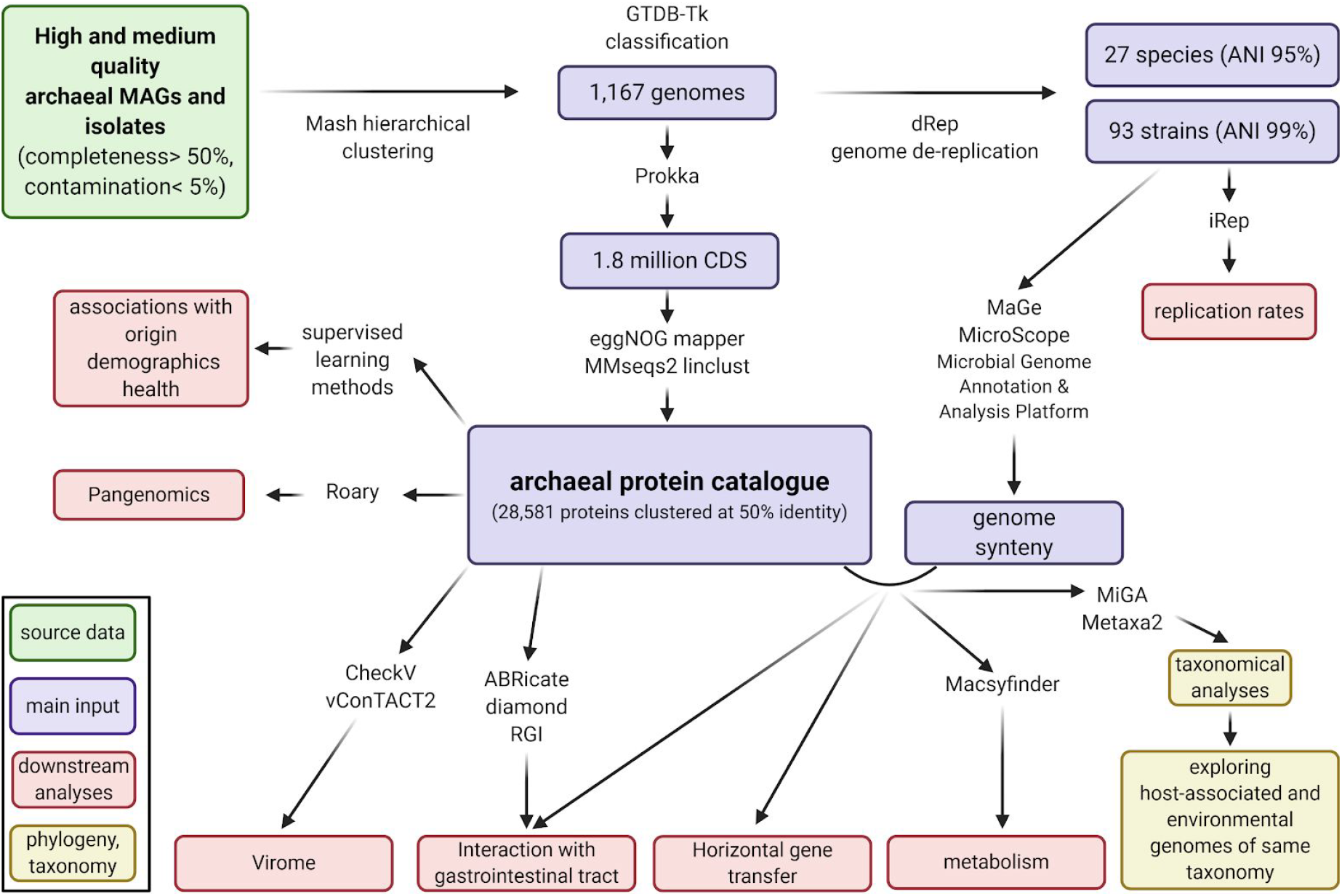
Methodology. Flow chart covering the major analysis steps of the study. Colored boxes show the source data (green), main input for the analysis (magenta), downstream analysis (red) and the phylogenetic and taxonomic analysis of the presented data set (yellow). Different steps are connected by arrows highlighting a selection of used bioinformatic tools for each step.

**Supplementary Fig. 2:**
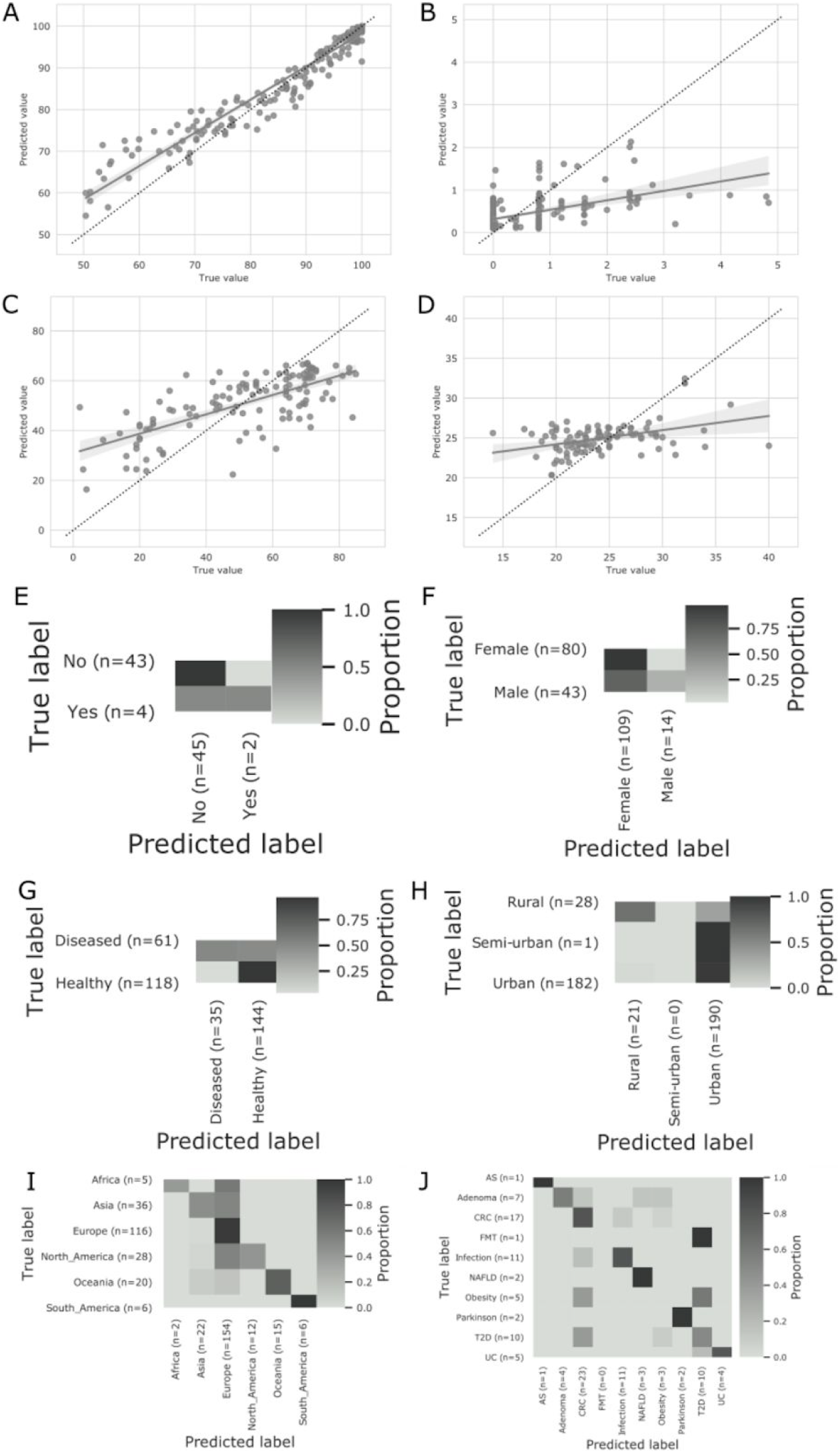
Predicting metadata values as a function of protein composition by supervised learning methods. Scatterplots and heatmaps of metadata predictions based on the unified archaeal MAG protein catalogue. Completeness (**A**), contamination (**B**), age (**C**), BMI (**D**), antibiotics (**E**), gender (**F**), health status (**G**), lifestyle (**H**), continent (**I**), and name of disease (**J**).

**Supplementary Fig. 3.**
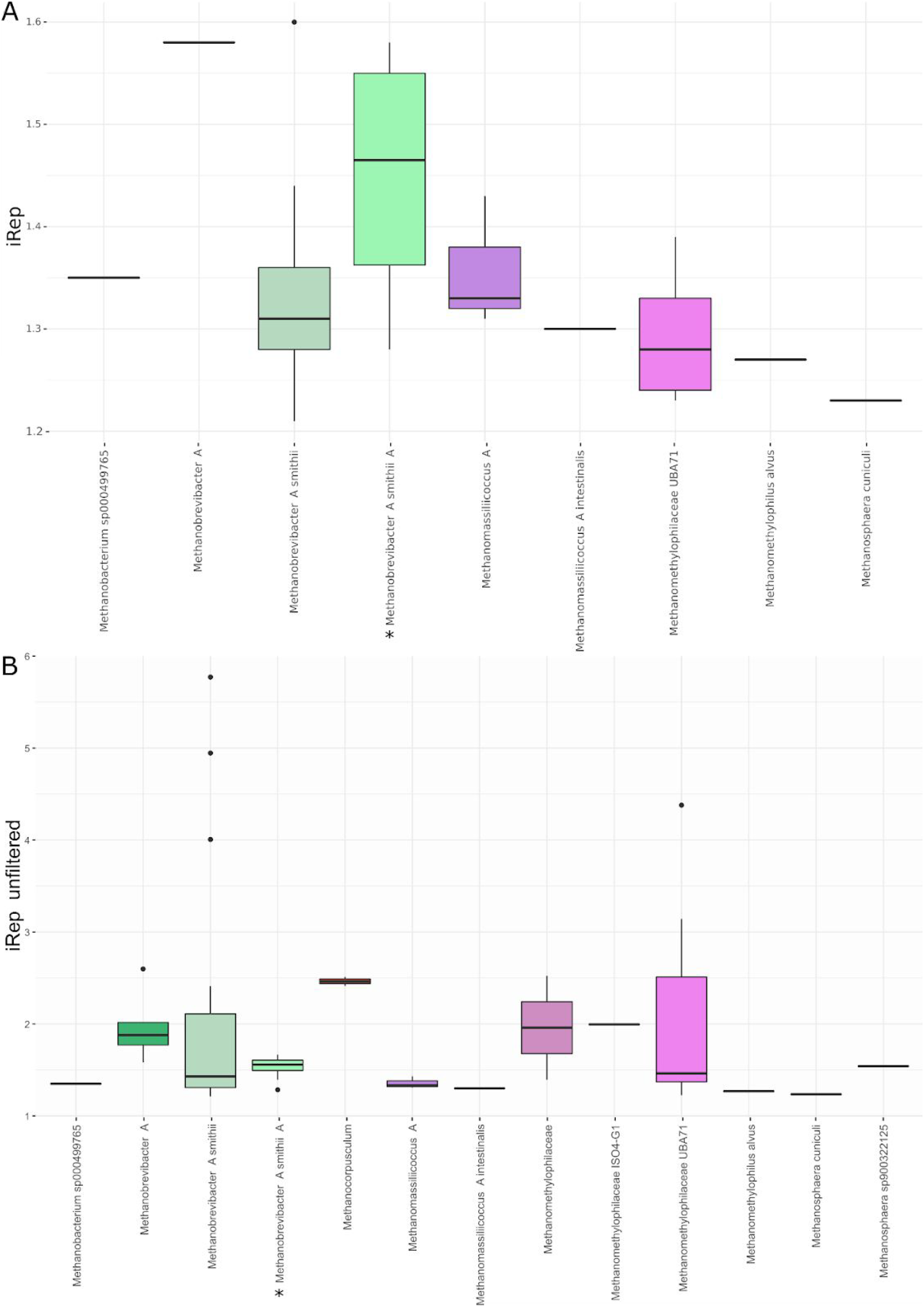
Replication rates of archaeal genomes from the human gut. **A**: iRep analysis of 26 genomes, showing the active replication of different GTDB-tk classified genomes. **B**: unfiltered iRep analysis on 69 genomes, showing the active replication of different GTDB-tk classified genomes. \**Candidatus* Methanobrevibacter intestini

**Supplementary Fig. 4:**
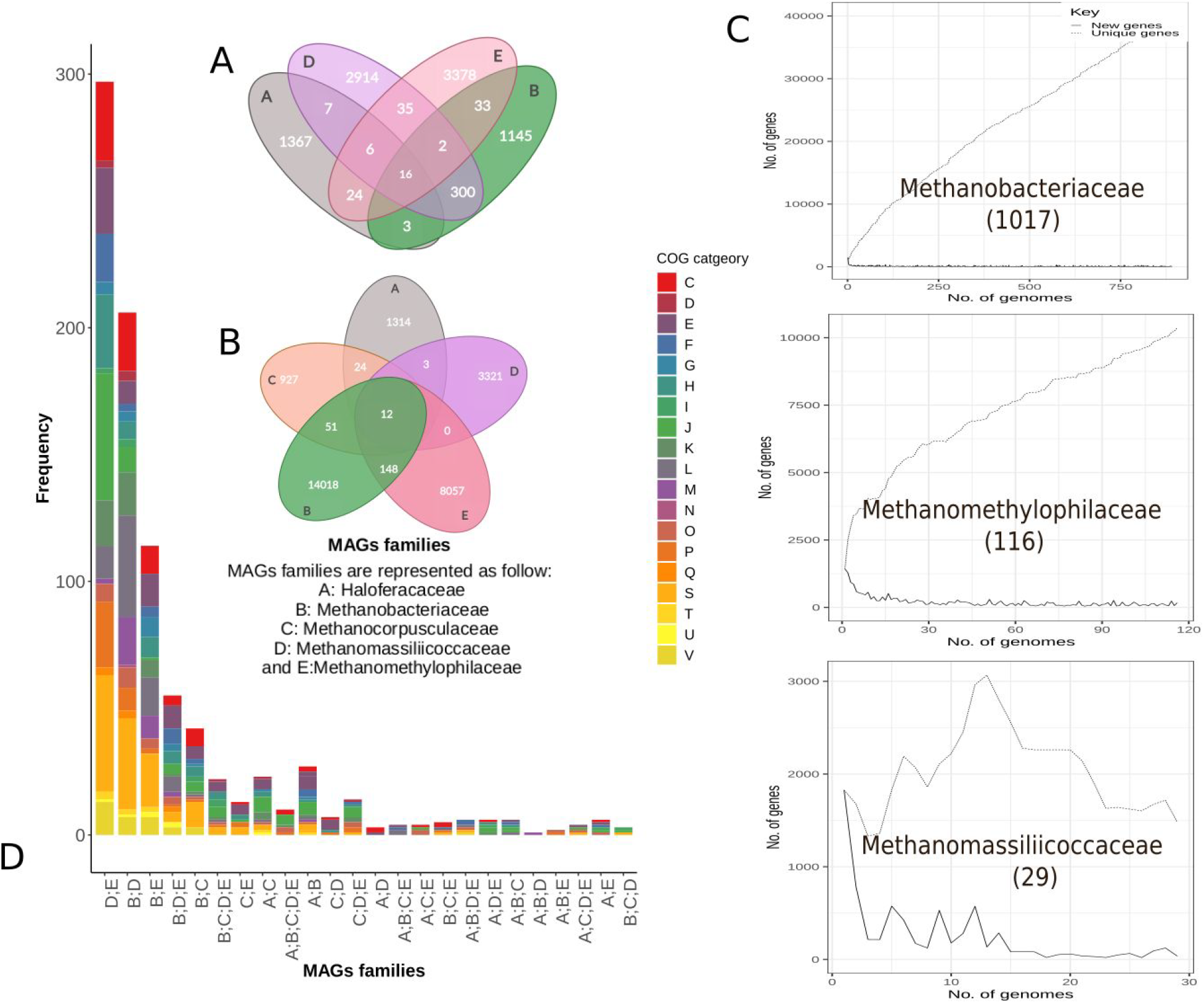
Venn diagrams summarizing the number of shared proteins between the different archaeal families. **A)** for complete genomes only **B)** for all 1167 genomes. Venn diagrams were done by creately (https://app.creately.com). **C)** Pan-genome analysis per family where over 10 members per family exist. The three archaeal families Methanomassiliicoccaceae (α=0.69), Methanobacteriaceae (α=0.71) and Methanomethylophilacea (α=0.86) have an open pangenome. **D)** Bar plot representing the frequency of COG categories of the proteins shared between the 5 archaeal MAGs taxonomic families (CELLULAR PROCESSES AND SIGNALING: [D] Cell cycle control, cell division, chromosome partitioning, [M] Cell wall/membrane/envelope biogenesis, [N] Cell motility, [O] Post-translational modification, protein turnover, and chaperones, [T] Signal transduction mechanisms, [U] Intracellular trafficking, secretion, and vesicular transport, [V] Defense mechanisms. INFORMATION STORAGE AND PROCESSING: [J] Translation, ribosomal structure and biogenesis, [K] Transcription, [L] Replication, recombination, and repair. METABOLISM: [C] Energy production and conversion, [E] Amino acid transport and metabolism, [F] Nucleotide transport and metabolism, [G] Carbohydrate transport and metabolism, [H] Coenzyme transport and metabolism, [I] Lipid transport and metabolism, [P] Inorganic ion transport and metabolism, [Q] Secondary metabolites biosynthesis, transport, and catabolism. POORLY CHARACTERIZED: [S] Function unknown) Supplementary Table 2.

**Supplementary Fig. 5:**
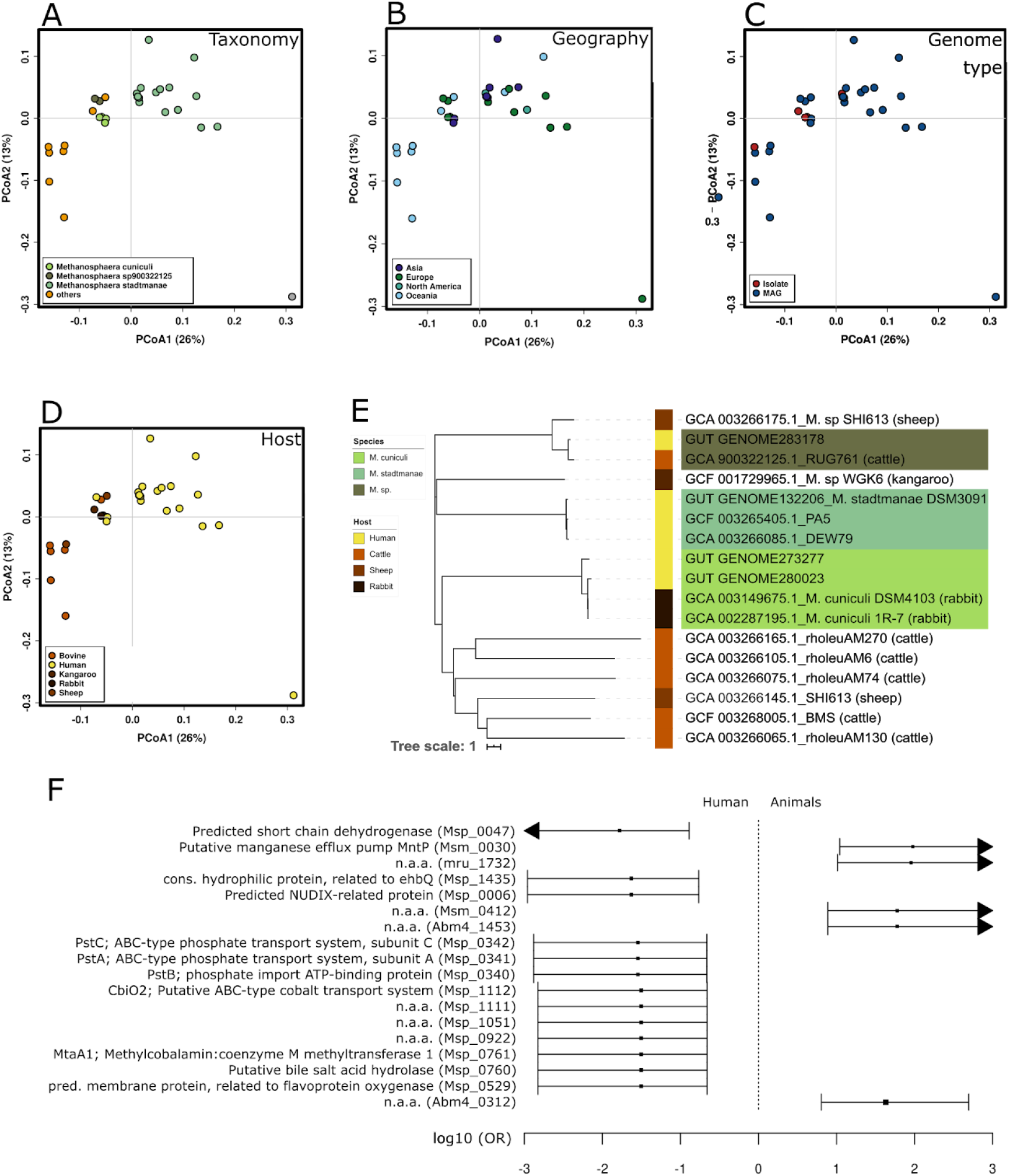
Profiles of human-associated *Methanosphaera* genomes. (Supplementary Table 9). For comparison, eleven genomes from animal-associated Methanosphaera were included. PCoA plots of the genomic profiles according to taxonomy (A), geography (B), genome type (C), and host (D) and phylogenetic tree of the genus *Methanosphaera* with human- and animal-associated representatives (E). Human-associated species are highlighted in green colors. Colored bar displays the origin: human (yellow) and animals (shades of brown). (F): Forest plot showing the outcome of the Wilcoxon rank test comparison of genomes from humans vs. animals (only proteins with FDR<0.05 are shown), bar displays the odds ratio (OR) (Supplementary Table 9).

**Supplementary Fig. 6:**
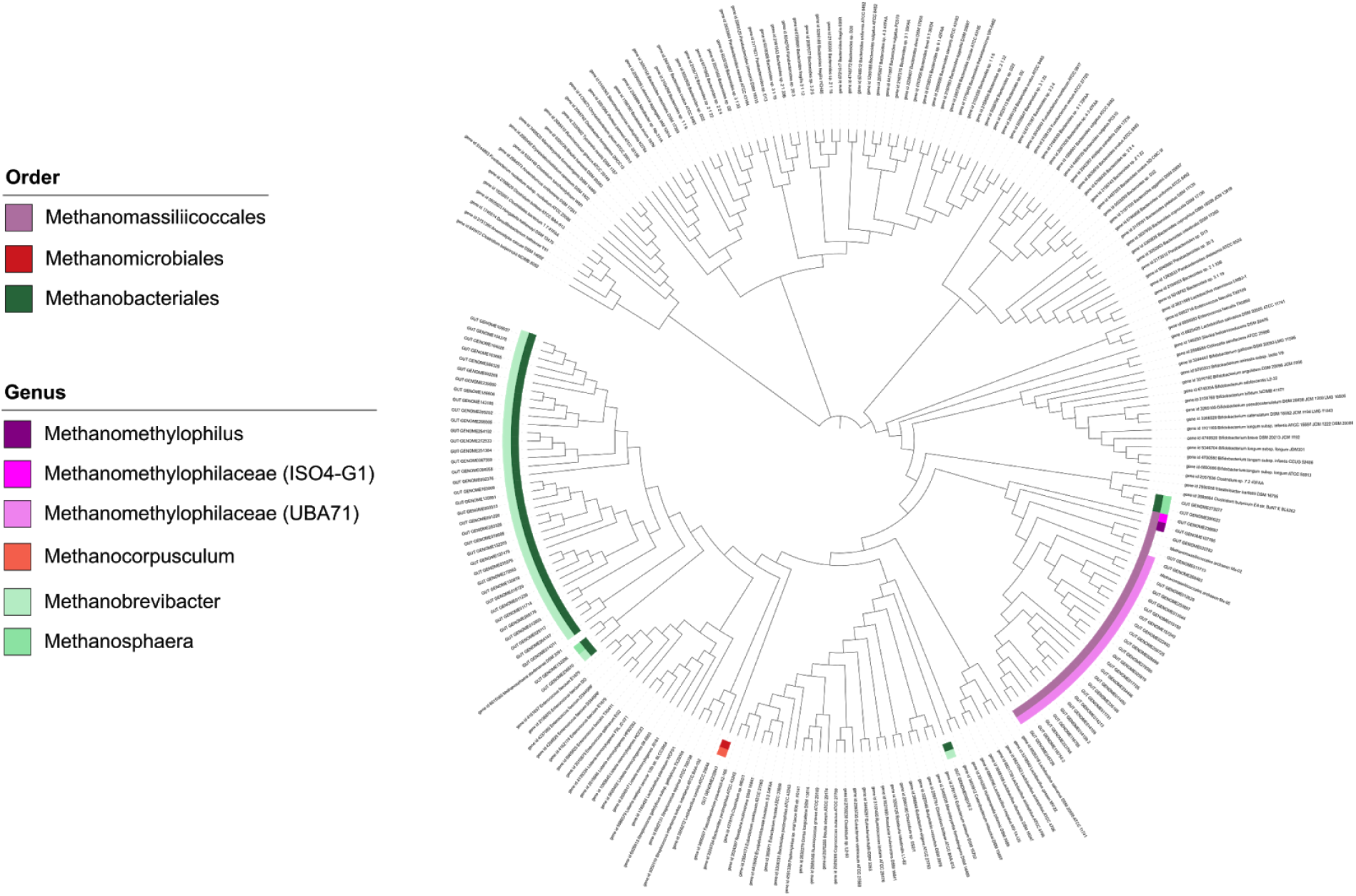
Archaeal bile salt hydrolase genes. (this study) integrated in the bacterial tree of BSHs (Song et al. 2019). Archaeal genes are highlighted by the colored ring, indicating the respective taxonomic affiliation.

**Supplementary Fig. 7:**
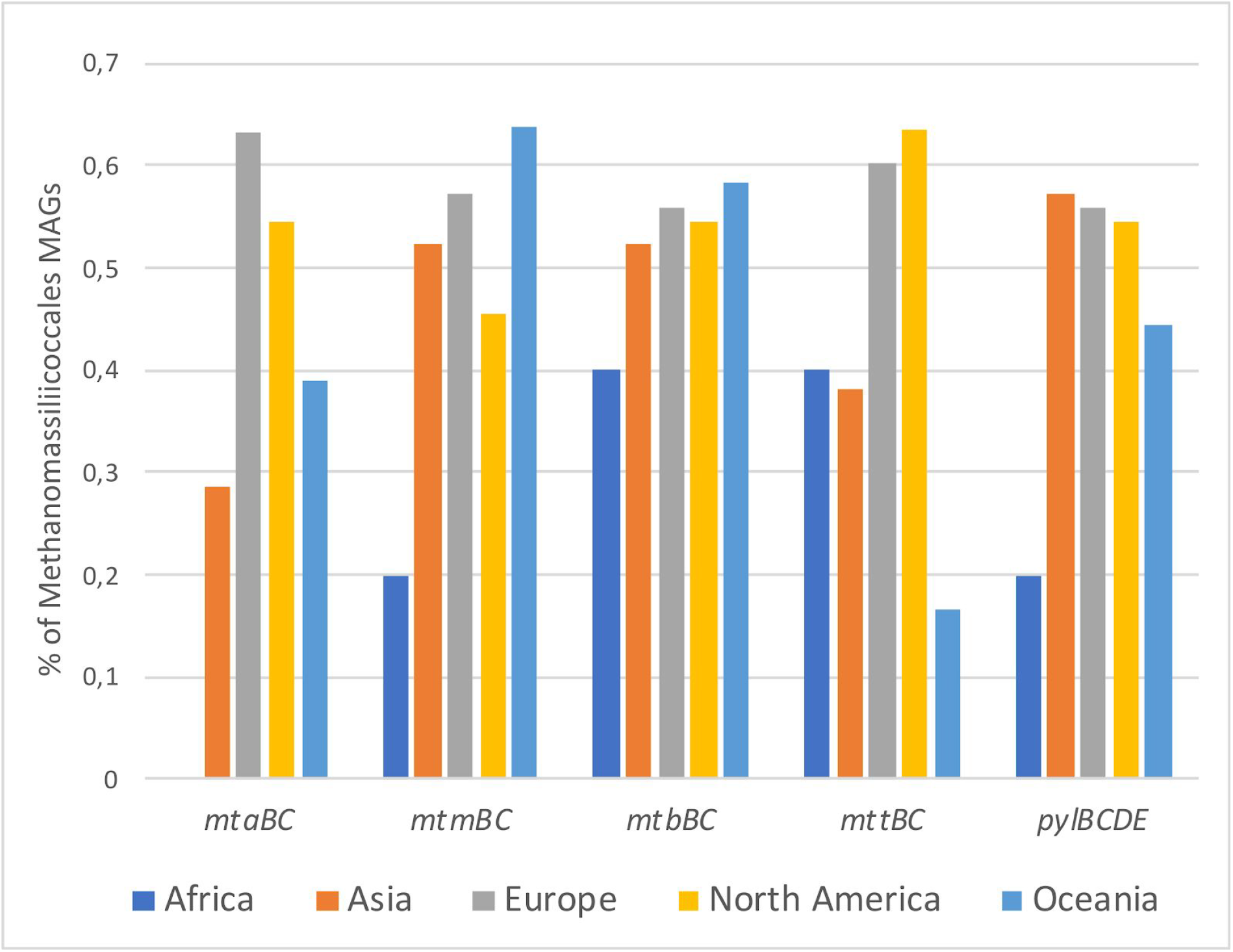
Geographic distribution of methyl-compound utilization capacity by Methanomassiliicoccales representatives. The presence of mtaBC, mtmBC, mtbBC and mttBC genes needed for methanol, monomethylamine, dimethylamine and, trimethylamine utilization, respectively, as well as pylBCDE genes responsible for the biosynthesis of pyrrolysine (an amino-acid specifically present in methylamine methyltransferases) was searched in all Methanomassiliicoccales MAGs. Methanomassiliicoccales were separated according to the geographic location (continents) of their host, and the percentage of them having the above mentionned genes is displayed.

**Supplementary Table 1.** Detailed description of all MAGs and isolates, which were analyzed in this study. This table includes the list of all 1167 archaeal genomes, a strain list (99% ANI distance), a species list (95% ANI distance) and all available metadata. These data sets were the basis for Fig. 1, Fig.3 and Supplementary Fig. 2,

**Supplementary Table 2.** Unified human archaeal protein catalogue based on clustering at 50% identity of all genome CDS, with associated lineage and genome information and a summary of the number of genes shared per archaeal family. Basis for Fig 3D and Supplementary Fig. 4 A,B and D.

**Supplementary Table 3.** Metadata information on the human and animal *Methanobrevibacter* analysis dataset, and protein catalogue information for those genomes (cut-off: presence in at least 10 genomes). Basis for Fig. 4.

**Supplementary Table 4.** Comparison of *M. smithii* vs. *M. smithii*_A, based on genome size and discriminative proteins (Wilcoxon top 25). Basis for information provided in chapter “The *Methanobrevibacter smithii* clade splits into two separate species”.

**Supplementary Table 5.** Identified proviruses and clustering into viral populations including quality summaries and host where they were identified from. Basis for Fig. 5.

**Supplementary Table 6.** Proportions of non-archaeal proteins for Methanosphaera, Methanobrevibacter and Methanomassiliicoccales, organized for human and animal-derived genomes, and isolates. Basis for Fig. 6.

**Supplementary Table 7. Comparison to environmental archaea. i)** Origin of 16S rRNA genes. Basis for Fig 7A. **ii)** Subset of the environmental archaeal genomes identified by Parks et al. 2017. Basis for Fig. 7 B,C,D and E.

**Supplementary Table 8.** Presence/absence of enzymatic complexes and pathways in the 27 species reported in this study. The presence/absence of the complexes and pathways is inferred from the detection of all enzymes that compose them. Three putative species represented by only 1 MAG having a poor quality were not included in this analysis. “+” indicates the presence of the complex or the pathways and “−” indicates its absence. MCR, Methyl-coenzyme M reductase; MTR, N5-Methyltetrahydromethanopterin: coenzyme M methyltransferase; H4MPT mWL, H4MPT version of the methyl branch of the Wood-Ljungdhal pathway; MtaBC, methanol:corrinoid protein methyltransferase and associated corrinoid-binding protein; MtmBC, monomethylamine:corrinoid protein methyltransferase and associated corrinoid-binding protein; MtbBC, dimethylamine:corrinoid protein methyltransferase and associated corrinoid-binding protein; MtbBC, trimethylamine:corrinoid protein methyltransferase and associated corrinoid-binding protein; MtsAB, Methylthiol:coenzyme M methyltransferase and associated corrinoid-binding protein; PylBCDE, pyl-system involved in pyrrolysine biosynthesis; FdhAB, formate dehydrogenase; "Alcohol utilisation" is based on the presence of Adh (alcohol dehydrogenase), Aldh (Aldehyde dehydrogenase) and Fno (NADPH-dependent F420 reductase); Fpo-like, Ferredoxin:heterodisulfide oxidoreductase; Eha, Energy-conserving hydrogenase A; Ehb, Energy-conserving hydrogenase B; Ech, Energy-conserving hydrogenase C; HdrBC, Heterodisulfide reductase catalytic subunit and C subunit; MvhADG, F420-nonreducing [Ni-Fe]-hydrogenase; FrhABG, F420H2-reducing hydrogenase; * in "Genome name" column indicates names of candidatus species and/or genera proposed in this study. Basis for Fig. 8B.

**Supplementary Table 9.** Metadata information on the human and animal *Methanosphaera* analysis dataset, protein catalogue information for those genomes, and Wilcoxon rank test comparison. Basis for Supplementary Fig. 5.

**Supplementary Material 1.** Alignment of the mcrA genes of M. smithii and M. smithii_A, fasta file. **Supplementary Material 2.** Environmental and human archaea 16S rRNA genes, fasta file, basis for Fig. 7A.

**Supplementary Material 3.** Interactive Krona output, basis for Fig. 6.

