## Supplementary Material 3 for "A comprehensive analysis of the global human gut archaeome from a thousand genome catalogue": Animal_Methanobrevibacter.html

Javascript must be enabled to view this page.

magnitude

 18533

 17415

 15083

 13551

 13551

 13551

 13551

 1427

 1427

 1427

 1427

 13

 13

 13

 13

 43

 43

 43

 43

 6

 6

 6

 6

 21

 21

 21

 21

 22

 22

 22

 22

 2326

 2326

 2326

 2326

 2326

 6

 6

 6

 6

 6

 0

 0

 0

 0

 0

 1108

 111

 111

 111

 111

 111

 651

 20

 20

 20

 20

 476

 340

 95

 11

 64

 3

 5

 2

 7

 2

 1

 68

 68

 114

 114

 30

 30

 13

 13

 19

 19

 1

 1

 0

 0

 131

 131

 131

 4

 4

 4

 1

 1

 1

 109

 36

 36

 36

 52

 25

 8

 16

 1

 0

 0

 0

 20

 20

 1

 1

 4

 4

 2

 2

 0

 0

 0

 0

 20

 13

 13

 2

 1

 1

 0

 0

 2

 2

 2

 2

 1

 1

 0

 0

 1

 1

 1

 3

 3

 3

 3

 11

 11

 11

 11

 32

 32

 32

 32

 117

 39

 23

 7

 7

 16

 16

 0

 0

 16

 16

 16

 9

 9

 9

 9

 10

 10

 10

 10

 18

 18

 18

 18

 41

 19

 19

 19

 22

 22

 14

 3

 3

 1

 1

 0

 0

 0

 0

 0

 0

 0

 0

 0

 18

 18

 18

 18

 18

 19

 19

 19

 19

 19

 3

 3

 3

 3

 3

 0

 0

 0

 0

 48

 48

 48

 48

 48

 76

 7

 7

 7

 7

 8

 1

 1

 1

 1

 1

 1

 1

 1

 1

 0

 0

 0

 0

 3

 3

 3

 1

 1

 1

 1

 1

 1

 35

 1

 1

 1

 7

 7

 7

 1

 1

 1

 15

 15

 15

 2

 2

 2

 6

 6

 6

 1

 1

 1

 1

 1

 1

 1

 1

 1

 0

 0

 0

 0

 0

 0

 0

 0

 0

 0

 0

 0

 0

 0

 0

 0

 0

 0

 0

 0

 0

 0

 0

 12

 11

 11

 11

 1

 1

 1

 0

 0

 0

 9

 9

 9

 9

 5

 2

 2

 2

 0

 0

 3

 3

 3

 0

 0

 0

 0

 0

 0

 0

 20

 7

 7

 7

 7

 7

 5

 3

 3

 2

 2

 2

 2

 2

 0

 0

 0

 6

 3

 3

 3

 3

 3

 3

 1

 1

 1

 1

 1

 3

 3

 3

 3

 3

 9

 9

 9

 9

 9

 18

 11

 11

 11

 11

 4

 4

 4

 4

 3

 2

 2

 2

 1

 1

 1

 0

 0

 0

 0

 0

 0

 0

 0

 0

 0

 0

 10

 10

 10

 10

 10

 1

 1

 1

 1

 1

 1

 1

 1

 1

 1

 2

 2

 2

 2

 2

 0

 0

 0

 0

 0

 0

 0

 0

 0

 0

 1

 1

 1

 1

 1

 1

 3

 2

 2

 2

 2

 2

 1

 1

 1

 1

 1

 0

 0

 0

 0

 0

 0

 0

 0

 0

 0

 0

 0

 0

 0

 6

 2

 2

 2

 2

 2

 4

 4

 4

 4

 4

 0

 0

 0

 0
