## Supplementary Material 3 for "A comprehensive analysis of the global human gut archaeome from a thousand genome catalogue": Animals_Methanosphaera.html

Javascript must be enabled to view this page.

magnitude

 9205

 8589

 7679

 7294

 7294

 7294

 7294

 340

 340

 340

 340

 14

 14

 14

 14

 12

 12

 12

 12

 10

 10

 10

 10

 7

 7

 7

 7

 2

 2

 2

 2

 905

 905

 905

 905

 905

 5

 5

 5

 5

 5

 612

 28

 2

 2

 2

 2

 7

 2

 2

 2

 0

 0

 0

 0

 0

 0

 4

 4

 4

 1

 1

 1

 4

 4

 4

 4

 2

 2

 2

 2

 3

 3

 3

 3

 0

 0

 0

 0

 9

 1

 1

 1

 7

 7

 7

 1

 1

 1

 1

 1

 1

 1

 40

 10

 8

 8

 8

 2

 2

 1

 1

 4

 4

 4

 4

 4

 4

 4

 4

 19

 5

 5

 5

 13

 11

 11

 1

 1

 1

 1

 1

 1

 1

 2

 2

 2

 2

 1

 1

 1

 1

 39

 39

 39

 39

 39

 464

 332

 240

 30

 30

 97

 97

 73

 14

 12

 31

 12

 3

 1

 0

 6

 6

 6

 6

 24

 24

 4

 4

 2

 2

 2

 80

 80

 80

 4

 4

 4

 6

 6

 6

 76

 20

 20

 20

 21

 10

 10

 5

 5

 1

 0

 1

 2

 2

 3

 3

 32

 11

 11

 17

 16

 0

 1

 0

 0

 3

 3

 1

 1

 3

 3

 3

 7

 7

 7

 7

 20

 20

 20

 20

 19

 19

 19

 19

 10

 10

 10

 10

 2

 2

 2

 2

 2

 6

 3

 3

 3

 3

 1

 1

 1

 1

 2

 2

 2

 2

 7

 7

 7

 7

 7

 3

 3

 3

 3

 3

 2

 2

 2

 2

 2

 13

 1

 1

 0

 0

 1

 1

 8

 8

 8

 8

 4

 4

 4

 4

 4

 4

 4

 4

 4

 3

 3

 3

 3

 3

 1

 1

 1

 1

 1

 4

 1

 1

 1

 1

 1

 3

 3

 3

 3

 3
