## Supplementary Material 3 for "A comprehensive analysis of the global human gut archaeome from a thousand genome catalogue": Human_Methanobrevibacter.html

Javascript must be enabled to view this page.

magnitude

 47776

 46404

 39169

 35847

 35847

 35847

 35847

 3261

 3261

 3261

 3261

 18

 18

 18

 18

 27

 27

 27

 27

 1

 1

 1

 1

 8

 8

 8

 8

 7

 7

 7

 7

 7229

 7229

 7229

 7229

 7229

 6

 6

 6

 6

 6

 0

 0

 0

 0

 0

 1360

 162

 162

 162

 162

 162

 798

 45

 45

 45

 45

 541

 391

 61

 8

 38

 0

 0

 1

 10

 1

 3

 119

 119

 150

 150

 38

 38

 11

 11

 9

 9

 1

 1

 2

 2

 147

 147

 147

 2

 2

 2

 1

 1

 1

 156

 51

 51

 51

 79

 58

 9

 45

 2

 1

 0

 1

 10

 10

 1

 1

 7

 7

 2

 2

 1

 1

 0

 0

 26

 8

 8

 3

 0

 1

 0

 2

 13

 13

 2

 2

 0

 0

 0

 0

 0

 0

 0

 8

 8

 8

 8

 8

 8

 8

 8

 40

 40

 40

 40

 168

 83

 42

 37

 37

 5

 5

 0

 0

 41

 41

 41

 4

 4

 4

 4

 7

 7

 7

 7

 37

 37

 37

 37

 37

 20

 20

 20

 17

 17

 12

 2

 0

 0

 0

 1

 1

 1

 0

 0

 0

 0

 0

 0

 24

 24

 24

 24

 24

 7

 7

 7

 7

 7

 5

 2

 2

 2

 2

 3

 3

 3

 3

 3

 3

 3

 3

 3

 116

 5

 5

 5

 5

 34

 3

 3

 3

 4

 4

 4

 26

 0

 0

 25

 25

 1

 1

 0

 0

 0

 0

 0

 0

 1

 1

 1

 59

 0

 0

 0

 1

 1

 1

 4

 4

 4

 27

 27

 27

 2

 2

 2

 3

 3

 3

 0

 0

 0

 7

 7

 7

 2

 0

 0

 1

 1

 1

 1

 4

 4

 4

 4

 1

 1

 1

 1

 2

 2

 3

 3

 3

 1

 1

 1

 1

 1

 1

 12

 10

 10

 10

 1

 1

 1

 1

 1

 1

 3

 3

 3

 3

 3

 1

 0

 0

 1

 1

 0

 0

 0

 2

 2

 2

 0

 0

 0

 0

 41

 32

 32

 32

 32

 5

 3

 2

 2

 1

 1

 1

 1

 1

 1

 1

 1

 4

 2

 2

 2

 2

 2

 2

 0

 0

 0

 0

 0

 6

 6

 6

 6

 6

 3

 3

 3

 3

 3

 18

 5

 5

 5

 5

 1

 1

 1

 1

 12

 0

 0

 0

 5

 5

 5

 2

 2

 2

 4

 3

 3

 1

 1

 1

 1

 1

 4

 4

 4

 4

 4

 2

 2

 2

 2

 2

 0

 0

 0

 0

 0

 2

 2

 2

 2

 2

 1

 1

 1

 1

 1

 0

 0

 0

 0

 0

 0

 0

 0

 0

 0

 0

 4

 0

 0

 0

 0

 0

 3

 1

 1

 1

 1

 2

 2

 2

 2

 0

 0

 0

 0

 0

 1

 1

 1

 1

 1

 8

 1

 1

 1

 1

 1

 7

 7

 7

 7

 7

 0

 0

 0

 0
