## Supplementary Material 3 for "A comprehensive analysis of the global human gut archaeome from a thousand genome catalogue": Humans_Methanosphaera.html

Javascript must be enabled to view this page.

magnitude

 9828

 9677

 8854

 8602

 8602

 8602

 8602

 247

 247

 247

 247

 0

 0

 0

 0

 3

 3

 3

 3

 2

 2

 2

 2

 0

 0

 0

 0

 0

 0

 0

 0

 823

 823

 823

 823

 823

 0

 0

 0

 0

 0

 151

 6

 0

 0

 0

 0

 5

 0

 0

 0

 1

 1

 1

 2

 2

 2

 2

 2

 2

 0

 0

 0

 0

 0

 0

 0

 0

 0

 0

 0

 0

 0

 0

 0

 1

 1

 1

 1

 0

 0

 0

 0

 0

 0

 0

 0

 0

 0

 0

 0

 0

 0

 11

 4

 4

 4

 4

 0

 0

 0

 0

 2

 2

 2

 2

 0

 0

 0

 0

 1

 0

 0

 0

 1

 1

 1

 0

 0

 0

 0

 0

 0

 0

 4

 4

 4

 4

 0

 0

 0

 0

 20

 20

 20

 20

 20

 109

 77

 65

 9

 9

 32

 32

 14

 2

 2

 4

 4

 1

 0

 1

 3

 3

 1

 1

 5

 5

 1

 1

 0

 0

 0

 11

 11

 11

 0

 0

 0

 1

 1

 1

 17

 3

 3

 3

 6

 0

 0

 3

 3

 2

 2

 0

 1

 1

 0

 0

 8

 1

 1

 3

 2

 1

 0

 2

 2

 1

 1

 1

 1

 0

 0

 0

 1

 1

 1

 1

 4

 4

 4

 4

 7

 7

 7

 7

 3

 3

 3

 3

 1

 1

 1

 1

 1

 0

 0

 0

 0

 0

 0

 0

 0

 0

 0

 0

 0

 0

 0

 0

 0

 0

 0

 0

 0

 0

 0

 0

 2

 2

 2

 2

 2

 2

 1

 1

 1

 1

 0

 0

 0

 0

 0

 0

 1

 1

 1

 1

 0

 0

 0

 0

 0

 0

 0

 0

 0

 0

 0

 0

 0

 0

 0

 0

 0

 0

 0

 0

 0

 0

 0

 0

 0

 0
