## Supplementary Material 3 for "A comprehensive analysis of the global human gut archaeome from a thousand genome catalogue": Methanomassiliicoccales.html

Javascript must be enabled to view this page.

magnitude

 29521

 22129

 19634

 14935

 14935

 14935

 14935

 2397

 2397

 2397

 2397

 1379

 1379

 1379

 1379

 241

 241

 241

 241

 193

 193

 193

 193

 190

 190

 190

 190

 180

 180

 180

 180

 119

 119

 119

 119

 2354

 2354

 2354

 2354

 2354

 101

 101

 101

 101

 101

 40

 40

 40

 40

 40

 7351

 4717

 3783

 940

 940

 940

 2608

 710

 710

 431

 431

 304

 304

 643

 242

 183

 88

 67

 34

 18

 9

 2

 229

 229

 159

 159

 68

 68

 24

 24

 23

 23

 17

 17

 178

 178

 178

 57

 57

 57

 494

 352

 170

 170

 151

 108

 9

 8

 8

 8

 8

 2

 10

 10

 7

 7

 4

 4

 4

 4

 4

 4

 2

 2

 93

 93

 93

 37

 18

 18

 9

 9

 6

 4

 2

 2

 2

 2

 2

 12

 12

 12

 158

 158

 158

 158

 163

 163

 163

 163

 104

 104

 104

 104

 15

 15

 15

 15

 619

 619

 619

 619

 619

 738

 326

 120

 120

 120

 102

 102

 102

 62

 62

 62

 28

 28

 28

 14

 14

 14

 186

 92

 92

 92

 22

 22

 22

 10

 10

 10

 10

 10

 10

 8

 8

 8

 8

 8

 8

 14

 8

 8

 4

 4

 2

 2

 6

 6

 6

 8

 6

 6

 2

 2

 2

 2

 2

 2

 2

 2

 2

 2

 2

 2

 105

 65

 65

 65

 14

 14

 14

 22

 12

 12

 6

 6

 2

 2

 2

 2

 2

 2

 2

 2

 2

 2

 44

 44

 44

 44

 17

 17

 17

 17

 48

 14

 14

 14

 10

 10

 10

 24

 10

 10

 8

 8

 6

 6

 10

 4

 4

 4

 4

 4

 4

 2

 2

 2

 2

 2

 2

 2

 301

 92

 92

 92

 92

 137

 78

 78

 78

 21

 9

 9

 6

 6

 6

 6

 10

 10

 8

 2

 8

 8

 8

 8

 4

 4

 4

 4

 3

 3

 3

 3

 3

 3

 2

 2

 2

 2

 2

 2

 2

 2

 2

 66

 66

 66

 66

 6

 6

 6

 6

 83

 83

 83

 83

 83

 362

 239

 64

 64

 64

 173

 56

 56

 111

 111

 6

 6

 2

 2

 2

 36

 36

 36

 36

 39

 14

 14

 14

 25

 21

 9

 8

 4

 4

 4

 15

 15

 15

 15

 14

 14

 14

 14

 11

 11

 11

 11

 8

 8

 8

 8

 59

 59

 59

 59

 59

 55

 55

 55

 55

 55

 37

 37

 37

 37

 37

 92

 38

 38

 38

 38

 30

 30

 30

 30

 16

 16

 16

 16

 8

 8

 8

 8

 36

 36

 36

 36

 36

 34

 34

 34

 34

 34

 29

 29

 29

 29

 29

 31

 24

 24

 24

 24

 7

 7

 7

 7

 66

 22

 22

 22

 22

 29

 25

 22

 22

 3

 3

 4

 4

 4

 15

 13

 13

 13

 2

 2

 2

 22

 22

 22

 22

 22

 18

 18

 18

 18

 18

 16

 16

 16

 16

 16

 22

 12

 12

 12

 12

 10

 10

 10

 10

 6

 6

 6

 6

 6

 4

 4

 4

 4

 4

 2

 2

 2

 2

 2

 2

 2

 2

 2

 2

 23

 12

 12

 12

 12

 12

 3

 2

 2

 2

 2

 1

 1

 1

 1

 6

 6

 4

 4

 4

 2

 2

 2

 2

 2

 2

 2

 2

 18

 12

 12

 12

 10

 10

 2

 2

 6

 6

 6

 6

 6
